# Harnessing endogenous repair mechanisms for targeted gene knock-in of bovine embryos

**DOI:** 10.1101/2020.06.09.143115

**Authors:** Joseph R. Owen, Sadie L. Hennig, Bret R. McNabb, Jason C. Lin, Amy E. Young, James D. Murray, Pablo J. Ross, Alison L. Van Eenennaam

## Abstract

Introducing useful traits into livestock breeding programs through gene knock-ins has proven challenging. Typically, targeted insertions have been performed in cell lines, followed by somatic cell nuclear transfer cloning, which can be inefficient. An alternative is to introduce genome editing reagents and a homologous recombination (HR) donor template into embryos to trigger homology-directed repair (HDR). However, the HR pathway is primarily restricted to actively dividing cells (S/G2-phase) and its efficiency is low in zygotes, especially for the introduction of large DNA sequences. The homology-mediated end joining (HMEJ)-based strategy harnesses HDR by direct injection of embryos, and has been shown to have an improved knock-in efficiency in non-dividing cells. The knock-in efficiency for a 1.8kb gene was contrasted when combining a gRNA/Cas9 ribonucleoprotein complex with either a traditional HR donor template, or a HMEJ template in bovine zygotes. The HMEJ template resulted in a significantly higher rate of gene knock-in as compared to the HR template (37.0% and 13.8%; P < 0.05). Additionally, more than a third of the knock-in embryos (36.9%) were non-mosaic. This approach will facilitate the one-step introduction of gene constructs at a specific location of the bovine genome and contribute to the next generation of elite cattle.

## INTRODUCTION

The use of genetics, selective breeding and reproductive technologies has had a positive impact on livestock genetic improvement^1^. However, there is still a need to further improve production traits in order to address food safety and security, as well as human health and nutrition^2^. One potential example of how recent advances in biotechnology could improve production traits is genome editing^3^. Genome editing has the potential to address concerns related to livestock health and welfare, as well as improving production traits^4^.

In livestock, the predominant method for creating a gene insertion or knock-in (KI) using genome editing uses a two-step process, in which first a primary cell line is modified in culture and then used for somatic cell nuclear transfer (SCNT) cloning to create an animal with the intended gene modification^5^. However, SCNT cloning is generally inefficient, and production of viable calves can be highly variable^6^. An alternative approach for increasing the efficiency of producing genome edited livestock is through direct cytoplasmic injection (CPI) of *in vitro* fertilized embryos^7^. However, the use of standard embryo culture and gene KI techniques by using CPI often results in embryos displaying high levels of genetic mosaicism^8^. We reported high rates of mosaicism when introducing CRISPR/Cas9 editing reagents into bovine embryos at 18 hours post insemination (hpi), especially when using Cas9 mRNA^9^. A recent study showed that introducing editing reagents into MII oocytes or early bovine zygotes (10hpi), prior to the first S-phase, reduced mosaicism^10^.

Many attempts have been made to increase the rate of homologous recombination (HR) or decrease the rate of non-homologous end joining (NHEJ) for gene insertion when using the CRISPR/Cas9 system via CPI of zygotes^11^. However, these approaches have been unsuccessful in bovine embryos as HR is primarily restricted to actively dividing cells^12^. Alternative homology directed repair (HDR) approaches have been utilized for KI using a donor template via the homology-mediated end joining (HMEJ) method^13^. This method has been shown to be active in gametes and early stage 1-cell embryos, in which proteins necessary for pushing DNA repair machinery towards the end-joining pathways are at their highest concentration^11^. While NHEJ utilizes the Ku proteins and a ligase to mend the double-strand break (DSB) by blunt end ligation, the HMEJ approach utilizes proteins for resection of the 5’ ends and annealing of homologous regions between the DSB and donor template similar to that in the microhomology-mediated end joining (MMEJ) pathway^14^.

In this study, we employed the HMEJ method for the precise insertion of the sex-determining region Y (*SRY*) gene into a region 10kb downstream of the zinc finger, X-linked (*ZFX*) gene on the X chromosome of bovine embryos by injecting either *in vitro* matured oocytes prior to *in vitro* fertilization, or presumptive zygotes 6hpi. We used two donor templates to compare KI efficiency using the HMEJ and HR approaches, and show an increased rate of gene insertion at the target location when using the HMEJ donor template.

## RESULTS

### Guide-RNA design and testing

Each of three guide-RNAs (gRNAs) targeting the ZFX locus were independently injected 6hpi alongside Cas9 protein into zygotes in groups of 30, including a non-injected group of 50 embryos as controls. Each treatment was repeated three times. The overall number of blastocysts and mutation rate was determined for each guide injected into zygotes (Supplemental Table S1). ZFXg3 showed a significantly higher mutation rate 81.8% compared to ZFXg1 and ZFXg2 (37.5% and 57.1%, respectively; P < 0.05). In addition, a significant decrease in development to the blastocyst stage was observed when comparing ZFXg2 and ZFXg3 injected embryos to control non-injected embryos (17.5% and 14.9 vs. 26.7%, respectively; P < 0.05). However, there was no significant difference between the development to the blastocyst stage when comparing ZFXg1 injected embryos to non-injected controls (25.8% vs. 26.7%; P > 0.05). Based on these data, ZFXg3 was selected for further embryo KI testing.

### Donor plasmid testing in bovine embryos

The HMEJ (hmejSRYp) and HR (hrSRYp) donor plasmids (Fig. 1) were each injected alongside ZFXg3 and Cas9 protein to determine KI efficiency in bovine embryos. The editing reagents were injected into 25 groups of 30 MII oocytes each 18 hours after maturation followed by *in vitro* fertilization, and into eight groups of 30 presumptive zygotes each 6hpi. A significant decrease in development to the blastocyst stage was observed when injecting the *SRY* donor plasmids in MII oocytes and zygotes 6hpi as compared to non-injected controls (11.0% and 17.6% vs. 29.6%, respectively; P < 0.01; Fig. 2a). In addition, there was a significant decrease in development to the blastocyst stage when injecting MII oocytes as compared to presumptive zygotes 6hpi (11.0% vs. 17.6%; P < 0.05).

**Figure 1.**
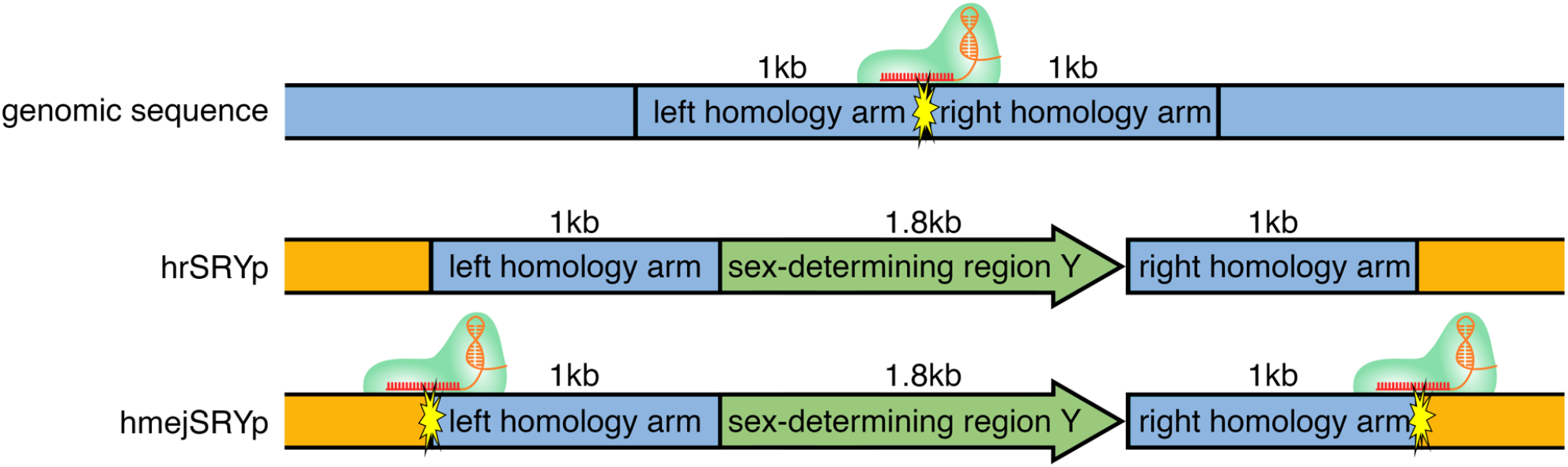
Schematic representation of donor plasmids used to test knock-in efficiency in bovine embryos. hrSRYp was used to compare the homologous recombination (HR) approach. hmejSRYp was used compare the homology mediated end-joining (HMEJ) approach. *SRY* = sex determining region Y; PAM = protospacer adjacent motif

**Figure 2.**
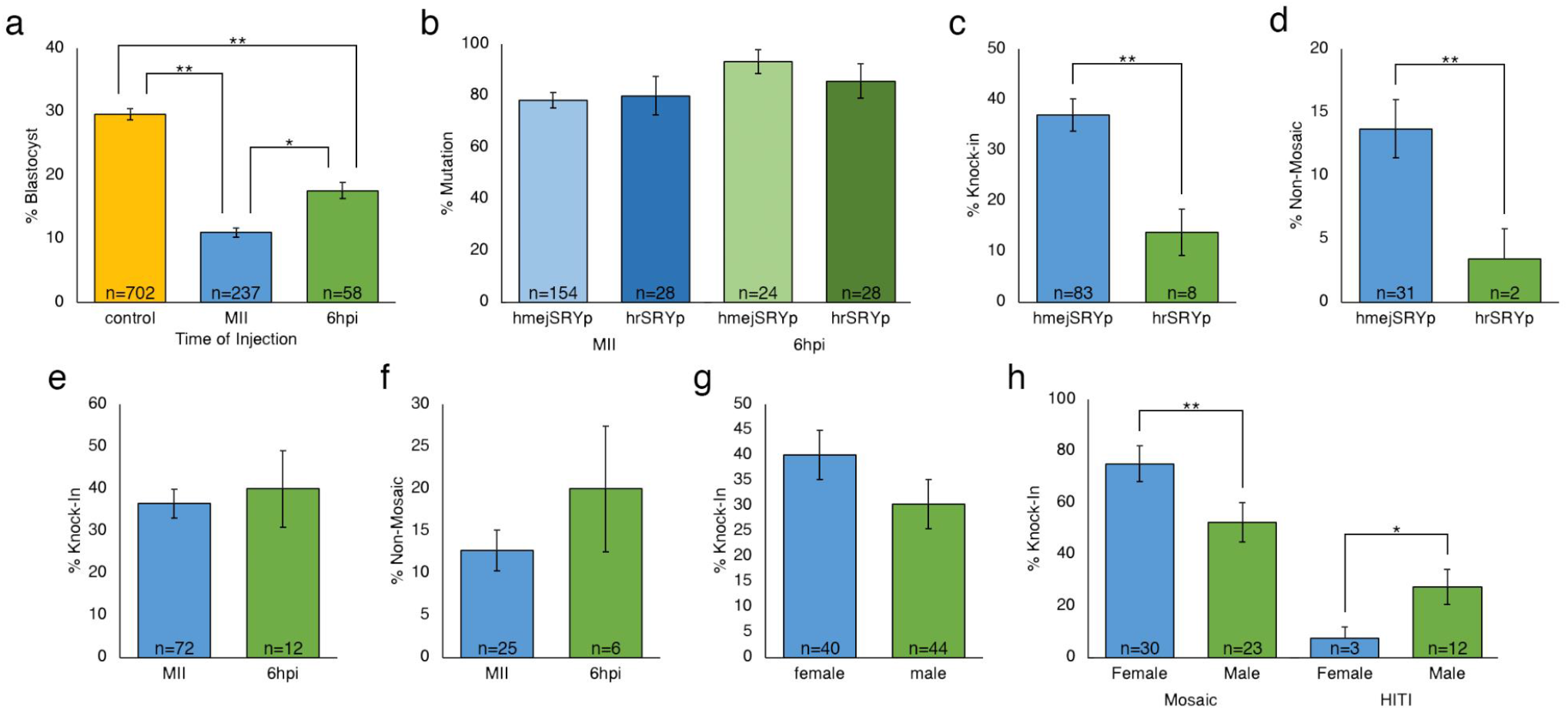
Comparison between development and knock-in efficiencies using hmejSRYp vs hrSRYp when injecting mature oocytes or presumptive zygotes six hours after insemination (6hpi). (a) Percentage of embryos that reached the blastocyst stage for oocytes injected prior to fertilization (MII) or presumptive zygotes injected 6hpi. (b) Percent mutations in embryos by donor template and time of injection. (c) Total percent knock-in embryos. (d) Percent non-mosaic knock-in embryos when comparing donor template. (e) Percent knock-in embryos when comparing time of injection. (f) Percent non-mosaic knock-in embryos when comparing time of injection. (g) Percent knock-in embryos when comparing sex of the embryo. (h) Evaluation of knock-in embryos by type of knock-in based on sex of the embryo. HITI = homology independent targeted insertion. Error bars = standard error of the mean. *P < 0.05; **P < 0.005.

Each blastocyst from the injected groups was analyzed for mutations induced at the target site, presence of KI and sex (Supplementary Information, Fig. S1). Overall mutation rates were not significantly different between male and female embryos, donor plasmids, or time of (P > 0.05; Table 1; Fig. 2b).

**Table 1.** Mutation, knock-in, and mosaicism rate in blastocysts after cytoplasmic injection of the gRNA:Cas9 RNP complex and hmejSRYp or hrSRYp in the MII oocyte or presumptive zygotes 6 hours post insemination. Homo = homozygous; Hemi = hemizygous. HITI = homology independent targeted insertion. Letters that differ in the same column are significantly different (P < 0.05).

| Sex | Time of Injection | Donor | n | % Mutation (n) | % Total Knock-In (n) | % Non-Mosaic Knock-In (n) | Subset of Knocked-In Embryos |  |  |
| --- | --- | --- | --- | --- | --- | --- | --- | --- | --- |
|  |  |  |  |  |  |  | Non-Mosaic |  | % Mosaic (n) |
|  |  |  |  |  |  |  | % Homo/Hemi (n) | % HITI (n) |  |
| Female | MII | hmejSRYp | 82 | 83 (68) | 42 (34)<sup>a</sup> | 10 (8)<sup>a</sup> | 18 (6)<sup>a</sup> | 6 (2)<sup>a</sup> | 76 (26)<sup>a</sup> |
|  |  | hrSRYp | 14 | 79 (11) | 29 (4)<sup>b</sup> | 14 (2)<sup>a</sup> | 50 (2)<sup>a</sup> | 0 (0)<sup>a</sup> | 50 (2)<sup>a</sup> |
|  | 6hpi | hmejSRYp | 13 | 92 (12) | 46 (6)<sup>a</sup> | 15 (2)<sup>a</sup> | 17 (1)<sup>a</sup> | 17 (1)<sup>a</sup> | 67 (4)<sup>a</sup> |
|  |  | hrSRYp | 16 | 88 (14) | 19 (3)<sup>b</sup> | 0 (0)<sup>a</sup> | 0 (0)<sup>a</sup> | 0 (0)<sup>a</sup> | 100 (3)<sup>a</sup> |
| Male | MII | hmejSRYp | 115 | 75 (86) | 33 (38)<sup>a</sup> | 15 (17)<sup>a</sup> | 21 (8)<sup>a</sup> | 24 (9)<sup>b</sup> | 55 (21)<sup>b</sup> |
|  |  | hrSRYp | 16 | 81 (13) | 6 (1)<sup>b</sup> | 0 (0)<sup>a</sup> | 0 (0)<sup>a</sup> | 0 (0)<sup>a</sup> | 100 (1)<sup>b</sup> |
|  | 6hpi | hmejSRYp | 17 | 94 (16) | 35 (6)<sup>a</sup> | 24 (4)<sup>a</sup> | 17 (1)<sup>a</sup> | 50 (3)<sup>b</sup> | 33 (2)<sup>b</sup> |
|  |  | hrSRYp | 12 | 83 (10) | 0 (0)<sup>b</sup> | n/a | n/a | n/a | n/a |
| Total | MII | hmejSRYp | 197 | 78 (154) | 37 (72)<sup>a</sup> | 13 (25)<sup>b</sup> | 19 (14) | 15 (11) | 66 (47) |
|  |  | hrSRYp | 30 | 80 (24) | 17 (5)<sup>b</sup> | 7 (2)<sup>a</sup> | 40 (2) | 0 (0) | 60 (3) |
|  | 6hpi | hmejSRYp | 30 | 93 (28) | 40 (12)<sup>a</sup> | 20 (6)<sup>b</sup> | 17 (2) | 33 (4) | 50 (6) |
|  |  | hrSRYp | 28 | 86 (24) | 11 (3)<sup>b</sup> | 0 (0)<sup>a</sup> | 0 (0) | 0 (0) | 100 (3) |

The KI efficiency was significantly increased (P < 0.01) for the hmejSRYp donor plasmid (37.0%) compared to the hrSRYp donor plasmid (13.8%), with 36.9% of the KIs being non-mosaic (Fig. 2c). The proportion of embryos that resulted in a non-mosaic KI blastocyst was significantly higher for hmejSRYp compared to hrSRYp (13.7% versus 3.4%, respectively; P < 0.01; Fig. 2d). However, there was no significant difference in KI efficiency when comparing blastocysts that were injected at the MII stage to presumptive zygotes injected 6hpi (36.5% and 40.0%, respectively; P > 0.05; Fig. 2e). In addition, there was no significant difference in the number of non-mosaic embryos when injecting the hmejSRYp donor plasmid at the MII stage compared to 6hpi (12.7% vs. 20.0%, respectively; Fig. 2f; p > 0.05).

Along with the presence of homozygous and hemizygous KI embryos resulting from HDR, there were 15 homozygous or hemizygous embryos that resulted from homology independent targeted insertion (HITI): 11 in MII injected oocytes (15.3%) and four in embryos injected 6hpi (33.3%; Table 1). Previous studies have shown insertion of a double-strand DNA product is possible by HITI^15^. Sequences were analyzed for the presence of the donor plasmid backbone that could integrate using the HITI approach, but no reads containing backbone sequences were found (Supplementary Information, Fig. S2). In addition, there was no alignment of the unsorted reads to the donor plasmid backbone.

There was no significant difference in KI efficiency based on sex of the embryo (40.0% female vs. 30.3% male; P > 0.05; Fig. 2g). However, there was a significant increase in the level of female mosaic KI embryos compared to male mosaic KI embryos (75.0% vs. 52.3%; P < 0.01; Fig. 2h). Additionally, there was an increased rate of blunt end ligation of the cleaved hmejSRYp donor plasmid by HITI-mediated KI in male embryos compared to female embryos (27.3% vs. 7.5%; P < 0.05; Fig. 2h).

### Evaluation of mosaicism

Blastocysts that contained the *SRY* KI based on PCR analysis (Supplementary Information, Fig. S1 lanes 1-4) from each of the four injected groups were evaluated for the level of mosaicism using PacBio sequencing (Fig. 3; Supplementary Information, Table S2). Overall, there was a 64.1% rate of mosaicism. There was no significant difference in the average number of alleles, proportion of wild type reads or the proportion of *SRY* KI reads between MII oocytes and presumptive zygotes injected with the hmejSRYp donor plasmid, and MII oocytes injected with the hrSRYp donor plasmid (Table 2). However, there was significant increase in the number of alleles and the proportion of wild type reads and a significant decrease in *SRY* KI reads in the three presumptive zygotes injected with the hrSRYp donor plasmid, compared to all other groups.

**Figure 3.**
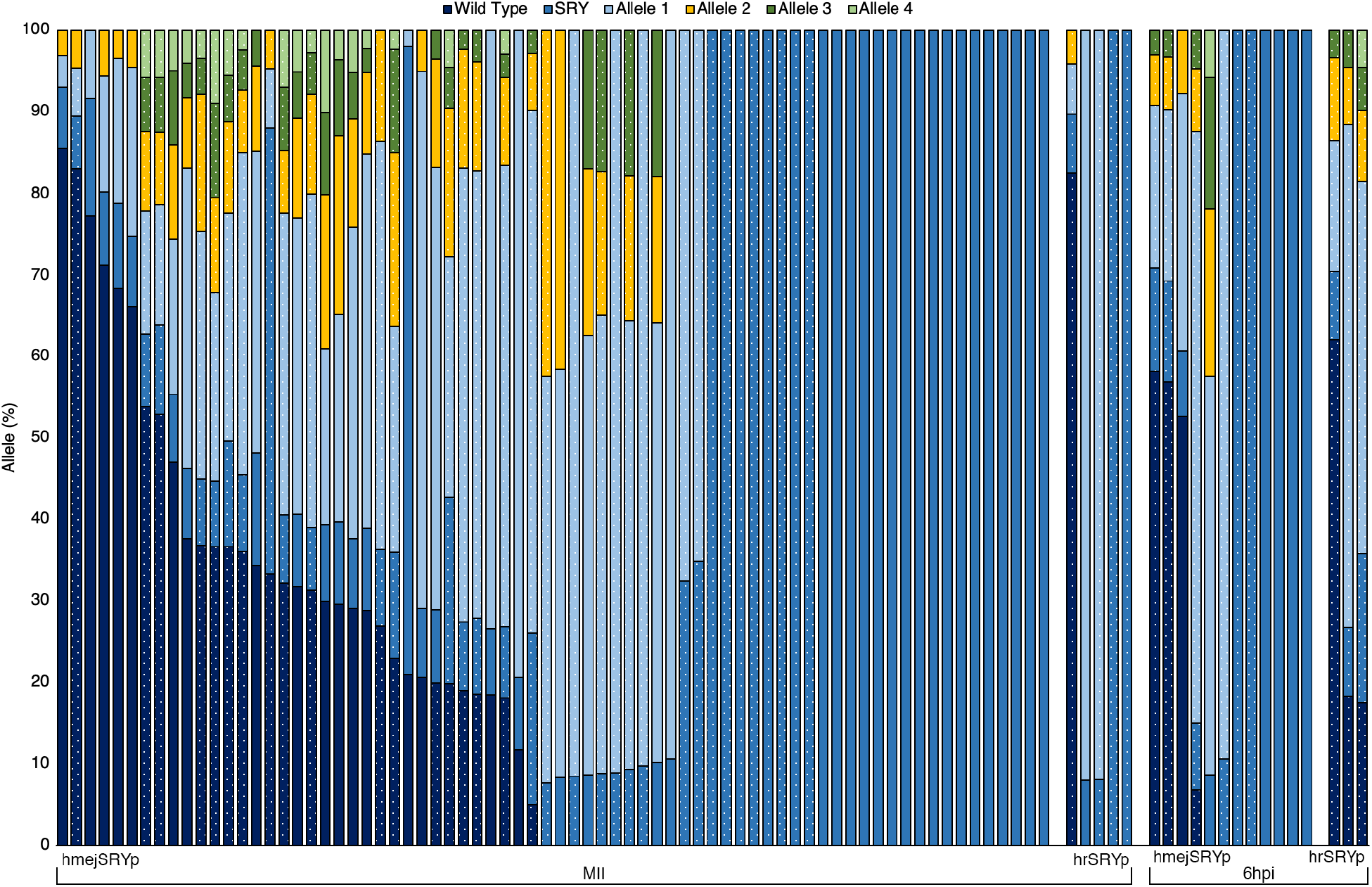
Bar graph depicting the percentage of alleles determined by PacBio sequencing in each of the 92 blastocysts that contained the SRY knock-in band in PCR analysis. Microinjection occurred either prior to fertilization (MII oocytes) or into presumptive zygotes 6 hours post insemination with either hmejSRYp or hrSRYp donor templates. dotted bars are female; solid bars are male.

**Table 2.** Average number of alleles, percent of wild type reads and percent of *SRY* knock-in reads by time of injection (MII oocytes or presumptive zygotes 6 hours post insemination) and donor template (hmejSRYp or hrSRYp). Letters that differ in the same column are significantly different (P < 0.05). SEM = standard error of the mean.

| Time | Donor | Alleles | SEM | % Wild Type | SEM | % SRY | SEM |
| --- | --- | --- | --- | --- | --- | --- | --- |
| MII | hmejSRYp | 3.3<sup>a</sup> | ±0.240 | 17.9<sup>a</sup> | ±2.8 | 43.4<sup>a</sup> | ±5.0 |
|  | hrSRYp | 2.0<sup>a</sup> | ±0.546 | 16.5<sup>a</sup> | ±16.5 | 44.7<sup>a</sup> | ±22.6 |
| 6hpi | hmejSRYp | 2.7<sup>a</sup> | ±0.554 | 14.5<sup>a</sup> | ±7.2 | 55.1<sup>a</sup> | ±13.6 |
|  | hrSRYp | 5.3<sup>b</sup> | ±0.335 | 32.6<sup>b</sup> | ±14.7 | 11.7<sup>b</sup> | ±3.3 |

Notably, within the group of 72 blastocysts derived from injected MII oocytes there were 35% (8 female and 17 male) non-mosaic KIs with the hmejSRYp donor plasmid (Table 2). Of these, two and nine were HITI, respectively. One of the female embryos contained roughly 50% *SRY* KI reads, suggesting a monoallelic KI. In addition, the wild type allele was also present, as well as two additional alleles containing indels for that sample, resulting in a mosaic blastocyst. Within this same group, two female embryos contained ~35% *SRY* KI reads with the remaining reads containing a ten base pair deletion at the CRISPR/Cas9 cut site. One of the male embryos contained ~75% *SRY* KI reads, suggesting that the insertion in one chromosome occurred after DNA replication in the zygote or in the 2-cell embryo, and that the other occurred after DNA replication at the 2-cell stage or at the 4-cell stage prior to DNA replication. In addition, two of the male embryos within this group contained ~25% *SRY* KI reads indicating that there was likely an insertion of *SRY* into one of the chromosomes after DNA replication at the 2-cell stage or at the 4-cell stage prior to DNA replication. The remaining mosaic samples across all four groups contained less than 20% *SRY* KI reads, with each of them containing more than three alleles.

### Embryo Transfers

Following analysis of blastocysts identifying the successful KI of *SRY* into the X chromosome, recipient animals were synchronized for embryo transfer of HMEJ donor injected MII oocytes followed by *in vitro* fertilization. For the first trial, a total of ten day-eight blastocysts were transferred to ten synchronized heifers (Supplementary Table S3). Embryo biopsies were taken prior to transfer and analyzed once the embryo transfer was complete. Three of the ten biopsies were positive for *SRY* insertion into the X chromosome. Two of the three were potential heterozygous/mosaic females and the third was a mosaic male. At day 35 of embryo development, pregnancies were diagnosed by transrectal ultrasonography, showing none of the recipients were positive for pregnancy. To limit the transfer of blastocysts that did not contain the *SRY* insertion, *SRY* KI blastocysts were produced over several rounds of MII oocyte injection, biopsied at the blastocyst stage and vitrified. A total of 12 *SRY* positive blasts were the used for embryo transfer. In addition, six non-biopsied fresh day-eight blastocysts produced from MII oocytes injected with the HMEJ donor were also transferred on the same day. Three of the recipients that received vitrified blastocysts and two of the six that received fresh day eight blastocysts were tentatively pregnant at day 35 based on ultrasound but were subsequently determined to be not pregnant at day 42.

## DISCUSSION

Using the HMEJ approach^13^ allowed for the targeted insertion of a large gene construct into the X-chromosome of bovine embryos. In 2017, Yao *et al.^13^,* showed this approach resulted in an increase in KI efficiency in mouse embryos to 22.7% compared to 11.9% when utilizing the MHEJ pathway, 3.3% when utilizing HR, and 1.4% when using HITI. Using the HMEJ approach, we observed a similar increase in KI efficiency (37.0%) compared to methods utilizing the traditional HR template (13.8%; ~2.7X increase). These results are similar to efficiencies previously reported in livestock species for KI via cytoplasmic injection of an HR template (5.7% in sheep embryos^16^ and 6.5% in pig embryos^17^). While there have been reports of using genome editors and donor template ODNs to induce single nucleotide polymorphisms or precise deletions of bovine embryos^18,19^, this is the first reported use of the CRISPR/Cas9 system for efficient KI of a large DNA segment (1.8kb).

To determine the optimal time for delivery of editing reagents to obtain non-mosaic KI embryos, we compared injection of the mature MII oocytes prior to fertilization to injection of zygotes 6hpi. Interestingly, a recent paper^10^ reported that 10hpi was the minimum bovine sperm and oocyte coincubation time required to obtain similar development rates to the conventional 20hpi IVF protocol, but we had acceptable rates of development with only 6 hours of gamete coincubation. The advantage of introducing editing reagents into the early zygote is the likelihood of reducing mosaicism by editing prior to the onset of DNA synthesis and the first cleavage division. Given that, we chose to introduce the editing reagents as early as possible (MII oocytes) and as late as feasible (6hpi) while still attempting to limit mosaicism. Injecting at these time points significantly reduced the number of embryos that reached the blastocyst stage compared to control non-injected embryos (11.0% and 17.6% compared to 29.6%). This decrease was likely a result of the DNA damage response illustrated by the decrease in the number of embryos that reached the blastocyst stage as the rate of mutation for a given guide increased. In addition, there was a further decrease in the number of embryos that reached the blastocyst stage when comparing injected MII oocytes (11.0%) to zygotes injected 6hpi (17.6%) due to the decrease in fertilization rate when subsequently co-culturing microinjected MII oocytes with cumulus-oocyte-complexes (COCs).

When comparing the KI efficiency between injected MII oocytes and zygotes injected 6hpi there was little difference observed (36.5% and 40.0%, respectively; p > 0.05). While the number of analyzed zygotes injected 6hpi is low; there were fewer embryos that reached the blastocyst stage when injecting MII oocytes compared to 6hpi. Therefore, it may be beneficial to inject 6hpi, as the KI efficiency is comparable but the latter yields roughly five percent more blastocysts per group. Likewise, a previous study found little difference in the editing efficiency when comparing the introduction of editing reagents into MII oocytes or 10hpi zygotes^10^. These authors suggested that the high degree of chromatin condensation in gametes may make it difficult for CRISPR-mediated editing in oocytes. This potentially indicates that the repair is occurring after fertilization, which would coincide with the similar KI efficiency observed when injecting MII oocytes and zygotes 6hpi.

The overall KI efficiency was higher using the HMEJ approach compared to the HR approach. Additionally, the rate of obtaining a non-mosaic embryo was also increased using the HMEJ approach. This increased rate of non-mosaic integration of *SRY* into the X chromosome (13.7%) is likely due to the DSB potentially being repaired by multiple repair mechanisms. This approach opens up the potential of HDR and MMEJ-mediated repair pathways, as well as HITI-mediated KIs. For the goal of inserting the 1.8kb *SRY* gene with promoter and coding sequence, the orientation of the insert is inconsequential, as transcription is not strand specific. However, in cases where precise changes, such as allelic substitutions or tagging a protein, are the ultimate goal, orientation of the insert is important. Using the HMEJ approach resulted in 7% of total injected embryos being non-mosaic KIs via an HDR-mediated pathway.

While HDR approaches have achieved successful KIs in cell lines, little success has been reported when using these approaches in livestock embryos. This is primarily due to the inactivity of the HR pathway in gametes and the early one-cell zygotes prior to the first round of DNA replication^11^ and the low efficiency of integration using the HITI approach^13^. The HMEJ approach utilized in this study has the ability to utilize the HDR pathway, the microhomology-mediated end joining (MMEJ) pathway and the NHEJ pathway, resulting in a HITI-mediated KI, which was observed here with 15 homozygous/hemizygous HITI-mediated KI embryos (6.6%)^13^. While the significant increase in mosaicism in female embryos can likely be attributed to the presence of two X chromosomes compared to one in the male embryos, the significant increase in HITI-mediated KI male embryos was unexpected. This may be explained by the uneven number of X chromosomes. As NHEJ is the primary mechanism for DNA repair in the early developing embryo, it is possible that the lack of a homologous mate resulted in the promotion of alternative pathways, driving the repair of the DSB by blunt end ligation.

The efficient integration of *SRY* into bovine embryos using the CRISPR/Cas9 approach has shown an overall improvement compared to previously reported studies. In a previous study, which characterized the level of mosaicism when introducing CRISPR/Cas9 editing reagents into bovine embryos 18hpi, we observed an average level of mosaicism of 94.2%^9^. Early delivery of CRISPR/Cas9 reagents combined with immediate and short activity of Cas9 can reduce mosaicism^20^. Here we found a decreased level of mosaicism after introducing CRISPR/Cas9 editing reagents into in MII oocytes (64.9%) and presumptive zygotes 6hpi (60.0%) (Table 2). This is not as low as the 30% reported when introducing targeted gene knock-outs in 6hpi zygotes^10^. However, in this study we only report mosaicism results for those samples with *SRY* present as we did not analyze mutations in samples with no *SRY* KI.

Similar, and sometimes higher, levels of mosaicism have been observed in studies with bovine, sheep and pigs compared to the level of mosaicism observed here^21–23^. Several strategies have been proposed to reduce mosaicism^20^ including altering the concentration of editing regents, the form of Cas9, and the time of injection, although no approach has eliminated mosaic mutations resulting from CRISPR/Cas9 genome editing. Ultimately, the efficiency of the gRNA and repair machinery to induce mutation prior to DNA replication in zygotes may be the most important factor to reduce mosaicism. The production of non-mosaic germline alterations is important in species like cattle as their two-year generation interval makes breeding mosaic founders to produce heterozygous or homozygous animals prohibitively expensive.

Although we were able to produce non-mosaic gene KI bovine embryos, the ultimate goal is to reliably and efficiently produce live animals. Here we have concluded further optimization needs to be undertaken to increase the efficiency of producing viable, non-mosaic gene KI embryos. *In vitro* produced embryos, as were used in this study, have reduced development potential relative to in vivo produced embryos^20^. Electroporation offers a promising approach to deliver editing reagents to zygotes without the physical damage of the microinjection technique ^24,25^ which may increase viability, although to date efficiencies of HDR editing in electroporated bovine embryos are low^26^.

The majority of KI animals created through electroporation of zygotes use single-stranded oligodeoxynucleotides (ssODN) donors, which have been shown to be stable and efficient in being incorporated into the genome through homology directed repair (HDR)^27^. Unfortunately, the size of the ssODN donor is a limitation as optimal ssODN donors are limited to a only about 100 base pairs ^28^, with the largest successful insertion using this approach being 1kb in length^29,30^.

Studies in mice have been able to overcome the size limitation of ssODN donors for embryo KI by using adeno-associated virus (AAV) to been successfully used to deliver the repair constructs in combination with CRISPR/Cas9 microinjection^31^ and to transfect large HDR donors of up to 4.9kb prior to electroporation^32^. Although this approach has yet to be applied to livesetock^30^, it has very high embryo survival rates with editing reported in up to 100% of offspring^33^.

Taking a biopsy from the trophectoderm of *in vitro* matured bovine embryos prior to transfer can result in live, healthy offspring^34^. However, a high level of skill is required to avoid decreasing embryo viability using this method and we were unsuccessful at obtaining pregnancies following embryo biopsy and vitrification. Others have used preimplantation diagnosis to successfully produce live HDR calves^19^, but even in that experiment only just over half of 24 high scoring cryopreserved embryos survived the process, and only three of the 13 transferred embryos established pregnancies. Another problem with preimplantation biopsies is mosaicism decreases the usefulness of these results^35^ as the trophectoderm may have a different genetic composition as compared to the inner cell mass. One potential alternative is the use of a non-invasive marker, such as a fluorescent protein, to screen for embryos with fluorescence in the inner cell mass prior to transfer, rather than an embryo biopsy. However, the presence of transgenic DNA in the resultant animal could pose a problem for food animal applications.

While there are still improvements to be made, this study explored an approach to facilitate the one-step introduction of a gene construct at a specific location in the bovine genome. Based on PCR using primers outside homology regions linking to the KI template, as well as utilizing PacBio sequencing, we demonstrated that KI embryos were generated, with the 1.8kb *SRY* gene construct correctly introduced into the target locus. Based on clonal sequences generated from PacBio, we could determine the presence and proportion of different alleles, determining the level of mosaicism. Taking this information into account we found that 36.9% of KI embryos were non-mosaic. This approach could be used to introduce novel genes or important genetic variants into livestock breeding programs.

## MATERIALS AND METHODS

### Animal Care

All experiments involving animals were approved and performed in compliance with the Institutional Animal Care and Use protocol #20595 at the University of California, Davis. Recipient cattle were maintained at the University of California, Davis Beef Barn.

### Embryo Production

Bovine ovaries were collected from a local slaughterhouse and transported to the laboratory at 35-37°C in sterile saline. Cumulus-oocyte-complexes (COCs) were aspirated from follicles and groups of 50 COCs were transferred to 4-well dishes containing 400μL of maturation media^36^. COCs were incubated for 18hr at 38.5°C in a humidified 5% CO_2_ incubator. Approximately 25 oocytes per drop were fertilized in 60uL drops of SOF-IVF^36^ with 1×10^6^ sperm per mL and incubated for 6hr at 38.5°C in a humidified 5% CO_2_ incubator. Presumptive zygotes were denuded by light vortex in SOF-HEPES medium^36^ for 5 min. 25 zygotes per drop were incubated in 50uL drops of KSOM culture media (Zenith Biotech, Glendale, CA, USA) at 38.5°C in a humidified atmosphere of 5% CO_2_, 5% O_2_, and 90% N_2_ for 7-8 days.

### Guide-RNA Design and Testing

Guide-RNAs were designed and tested as previous described^9^. In short, guides sequences were designed targeting the X-chromosome, 10kb downstream of the *ZFX* gene, with no less than three mismatches in the guide sequence for off-target sites and at least one mismatch in the seed region (8-11bp upstream of the PAM sequence) when compared to the bovine reference genome. Mutation rate for each guide was determined by laser-assisted cytoplasmic injection^37^ of *in vitro* fertilized embryos with 6pL of a solution containing 67ng/μL of *in vitro* transcribed gRNA alongside 133ng/μL of Cas9 mRNA or 167ng/μL of Cas9 protein (PNA Bio, Inc., Newbury Park, CA) incubated at room temperature for 30 minutes prior to injection. Embryos that reached blastocyst stage were lysed in 10μL of Epicenter DNA extraction buffer (Lucigen, Palo Alto, CA) using a SimpliAmp Thermal Cycler (Applied Biosystems, Foster City, California) at 65°C for 6 minutes, 98°C for 2 minutes and held at 4°C. The target region was amplified by two rounds of the polymerase chain reaction (PCR) using primers developed using Primer3 (Supplementary Information, Table S4)^38,39^. The first round of PCR was performed on a SimpliAmp Thermal Cycler (Applied Biosystems, Foster City, California) with 10μL GoTaq Green Master Mix (Promega Biosciences LLC, San Luis Obispo, CA), 0.4μL of each primer at 10mM and 9.2μL of DNA in lysis buffer for 5 min at 95°C, 35 cycles of 30 sec at 95°C, 30 sec at anneal temp (Supplementary Information, Table S4), and 30 sec at 72°C, followed by 5 min at 72°C. The second round of PCR was run with 10μL GoTaq Green Master Mix (Promega Biosciences LLC, San Luis Obispo, CA), 4.2μL of water, 0.4μL of each primer at 10mM and 5μL of first round PCR for 3 min at 95°C, 35 cycles of 30 sec at 95°C, 30 sec at anneal temp (Supplementary Information, Table S4), and 30 sec at 72°C, followed by 5 min at 72°C. Products were visualized on a 1% agarose gel using a ChemiDoc-ItTS2 Imager (UVP, LLC, Upland, CA), purified using the QIAquick Gel Extraction Kit (Qiagen, Inc., Valencia, CA) and Sanger sequenced (GeneWiz, South Plainfield, NJ).

### Donor Plasmid Construction

Donor plasmids were created to introduce *SRY* into the ZFX locus on the X-chromosome 10kb downstream of the *ZFX* gene. The donor plasmids were synthesized by GeneWiz (South Plainfield, NJ) to contain the endogenous *Bos taurus SRY* promoter and coding sequence (Accession U15569)^40^. 1kb homology arms were synthesized by GeneWiz containing regions flanking the cut site (South Plainfield, NJ) and inserted into the donor plasmids using Gibson Assembly Master Mix (New England Biolabs, Inc., Ipswich, MA), with (hmejSRYp) and without (hrSRYp) the endogenous CRISPR target site flanking the homology arms (Fig. 2). Plasmids were clonally amplified using 5-alpha Chemically Competent *E. coli* (High Efficiency) (New England Biolabs, Inc., Ipswich, MA) and extracted using the EndoFree Plasmid Maxi Kit (Qiagen, Inc., Valencia, CA).

### Cytoplasmic Injection and PCR Amplification for KI Efficiency

Knock-in of donor plasmids was attempted using laser-assisted cytoplasmic injection^37^ of *in vitro* matured oocytes after 18 hours of maturation and *in vitro* fertilized embryos 6 hours post insemination (hpi) with 6pL of a solution containing 67ng/μL of *in vitro* transcribed gRNA, 167ng/μL of Cas9 protein (PNA Bio, Inc., Newbury Park, CA) and 133ng/μL of donor plasmid. Injected MII oocytes were subsequently co-cultured with cumulus-oocyte-complexes (COCs) and *in vitro* fertilized following procedures previously described for sheep^22^. Embryos were scored for developmental stage reached at day 7-8. Embryos that reached blastocyst stage were lysed as described above and underwent whole-genome amplification using the Repli-G Mini kit (Qiagen, Inc., Valencia, CA). Target regions were amplified using primers developed using Primer3 (Supplementary Information, Fig. S1 and Table S4)^38,39^. PCR was performed on a SimpliAmp Thermal Cycler (Applied Biosystems, Foster City, California) with 12.5μL LongAmp *Taq* 2X Master Mix (New England Biolabs, Inc., Ipswich, MA), 9.5μL of H2O, 1μL of each primer at 10mM and 1μL of DNA for 5 min at 94°C, 35 cycles of 30 sec at 94°C, 30 sec at anneal temp (Supplementary Information, Table S4) and 4 min at 65°C, followed by 15 min at 65°C. Products were visualized on a 1% agarose gel using a ChemiDoc-ItTS2 Imager (UVP, LLC, Upland, CA), purified using the QIAquick Gel Extraction Kit (Qiagen, Inc., Valencia, CA) and Sanger sequenced (GeneWiz, South Plainfield, NJ).

### Next-Generation Sequencing of Knock-In Samples

Samples that were positive for *SRY* knock-in were PCR amplified using a dual round PCR approach described above to barcode samples for pooled sequencing (Supplemental File S2). Barcoded amplicons underwent SMRTbell library preparation and were sequenced on a PacBio Sequel II sequencer by GENEWIZ, LLC (South Plainfield, NJ, USA). Consensus sequences were called, reads separated by barcode and BAM converted to individual FASTQ files using SMRT Link v8.0.0.80529 (https://www.pacb.com/support/software-downloads/). Raw reads were aligned to each target site using Bowtie2-default v2.3.4.1^41^ (Supplementary Information, Fig. S2). Alignments were visualized using Integrative Genomics Viewer v2.6.2^42^. For mosaic analysis, reads were aligned to each target site using BWA v0.7.16a^43^. SAM files were converted to BAM files, sorted and indexed using SAMtools v1.9^44^. Number and types of alleles were determined for each sample using CrispRVariants v1.12.0^45^.

### Embryo Biopsy and Vitrification

Biopsies were performed seven days post-fertilization on all embryos that reached the blastocyst stage as previously described^35^. In short, small 8-10 cell biopsies were taken from the trophectoderm of expanded blastocysts using a microblade in Dulbecco’s phosphate-buffered saline lacking magnesium chloride or calcium chloride. Biopsies were transferred to 10μL of Epicenter DNA extraction buffer (Lucigen, Palo Alto, CA) and lysed as described above. Blastocysts were returned to culture media for two hours to monitor re-expansion. Re-expanded blastocysts were non-surgically transferred to synchronized recipients, as described below, or vitrified for transfer at a later date. Blastocysts were vitrified using Vit Kit-Freeze (Irvine Scientific, Santa Ana, CA, USA). Embryos were placed in ES solution until re-expanded, followed by 2 min in VS solution before being transferred to a CryoTip straw (Irvine Scientific, Santa Ana, CA, USA). The straw was the sealed and plunged into liquid nitrogen prior to transfer to a Dewar for storage. Embryos were thawed for transfer using Vit Kit-Thaw (Irvine Scientific, Santa Ana, CA, USA). Straws were thawed in a 37°C water bath for 3 sec, followed by transfer of the embryo to TS solution for 1 min. The embryo was transferred to DS solution for 4 min, then washed twice in WS solution prior to loading the embryo into the straw for transfer.

### Embryo Transfer

Recipient estrus synchronization was initiated by inserting an intravaginal progesterone device (1.38g; Eazi-Breed CIDR; Zoetis) and intramuscular administration of gonadotropin (100mcg; Factrel; Zoetis) on day 0 (sixteen days prior to transfer). On day 7, the CIDR was removed and intramuscular prostaglandin (25mg; Lutalyse; Zoetis) was administered. Recipients were monitored for estrus, and a second intramuscular dose of gonadotropin (100mcg; Factrel; Zoetis) was administered on day 9. Prior to transfer on day 16, recipient response to synchronization was confirmed via detection of an appropriate corpus luteum with transrectal ultrasonography. Prior to transfer, each recipient received a caudal epidural using 100mg 2% lidocaine (Xylocaine; Fresenius). Embryos were transferred via non-surgical, transcervical technique, and the blastocyst was deposited into the uterine horn ipsilateral to the corpus luteum. Pregnancy was diagnosed on day 35 of embryonic development by transrectal ultrasonography (5.0 MHz linear probe; EVO Ibex, E.I. Medical Imaging).

### Statistical Analysis

Comparison between development for guide analysis and mutation rates were evaluated using a linear model and statistical significance was determined using a Chi-square test. To analyze the level of mosaicism, an ANOVA test was used to determine significance between number of alleles per sample when injecting MII oocytes compared to 6hpi. For knock-in evaluation, injected groups or non-injected controls were blocked by replicate to determine the significance between the means in R using the Tukey method. A generalized linear model was run for samples to determine the effects of sex, timing and donor plasmid on mutation rates and knock-in efficiency. Each blastocyst was considered an individual test, evaluated using a binomial distribution and analyzed using a linear regression model. Once the unimportant variables were excluded, the groups were blocked by type and a two by two Chi square test was used to determine significance. Differences were considered significant when P < 0.05.

## Acknowledgements

The authors would sincerely like to thank Donald Harper, Marissa Fisher and Danny Vickers for excellent management of the animal herds, Ashley Young and Rebecca Ozeran for diligent collection and transport of ovaries from the processing plant to UC Davis and Erika Paulson for assisting with embryo injections. This project was supported by Biotechnology Risk Assessment Grant Program competitive grant no. 2015-33522-24106 from the U.S. Department of Agriculture, the Academic Federation Innovation Development Award at UC Davis, the Russell L. Rustici Rangeland and Cattle Research Endowment in the College of Agricultural and Environmental Science at UC Davis, the California Agricultural Experiment Station of the University of California, Davis, the Henry A. Jastro Research Fellowship in the College of Agricultural and Environmental Science at UC Davis and the National Institute for Food and Agriculture National Needs Graduate and Postgraduate Fellowship no. 2017-38420-26790 from the U.S. Department of Agriculture.

## Author’s contributions

JRO and SL performed the experiments with additional input from PJR, JDM and ALV. JRO, SLH, JCL, and AEY participated in sample processing and data analysis. BRM was responsible for embryo transfers and monitoring of pregnancies. JRO, SLH, JDM and ALV wrote the manuscript with suggestions from all the co-authors. All authors read and approved the final version.

## Data Availability

Raw sequence reads from PacBio Sequel II and Illumina MiSeq sequencing are available in the NCBI Sequence Read Archive as BioProject PRJNA635115 and SRA accession number SRR11850981. Individual results for the blastocyst development and mutation rate from each replicate (~ 30 embryos) of control and microinjected embryos are available in Supplementary Table S5-6.

## Supplemental Information

**Supplementary Table S1.** Mutation rate in embryos for each guide analyzed using PCR and Sanger sequencing. Multiple guides were tested targeting each locus to obtain highest efficiency guide. Letters that differ in the same column are significantly different (P < 0.05). Each chromosome independently tested using a two-by-two χ^2^ test.

| Allele | gRNA | Injected Embryos | % Blastocysts (n) | Blastocysts Analyzed | % Mutation Rate (n) |
| --- | --- | --- | --- | --- | --- |
| ZFX | 1 | 62 | 26 (16)<sup>a</sup> | 16 | 38 (6)<sup>a</sup> |
|  | 2 | 80 | 18 (14)<sup>b</sup> | 14 | 57 (8)<sup>a</sup> |
|  | 3 | 94 | 15 (14)<sup>b</sup> | 11 | 82 (9)<sup>b</sup> |

**Supplementary Table S2.**
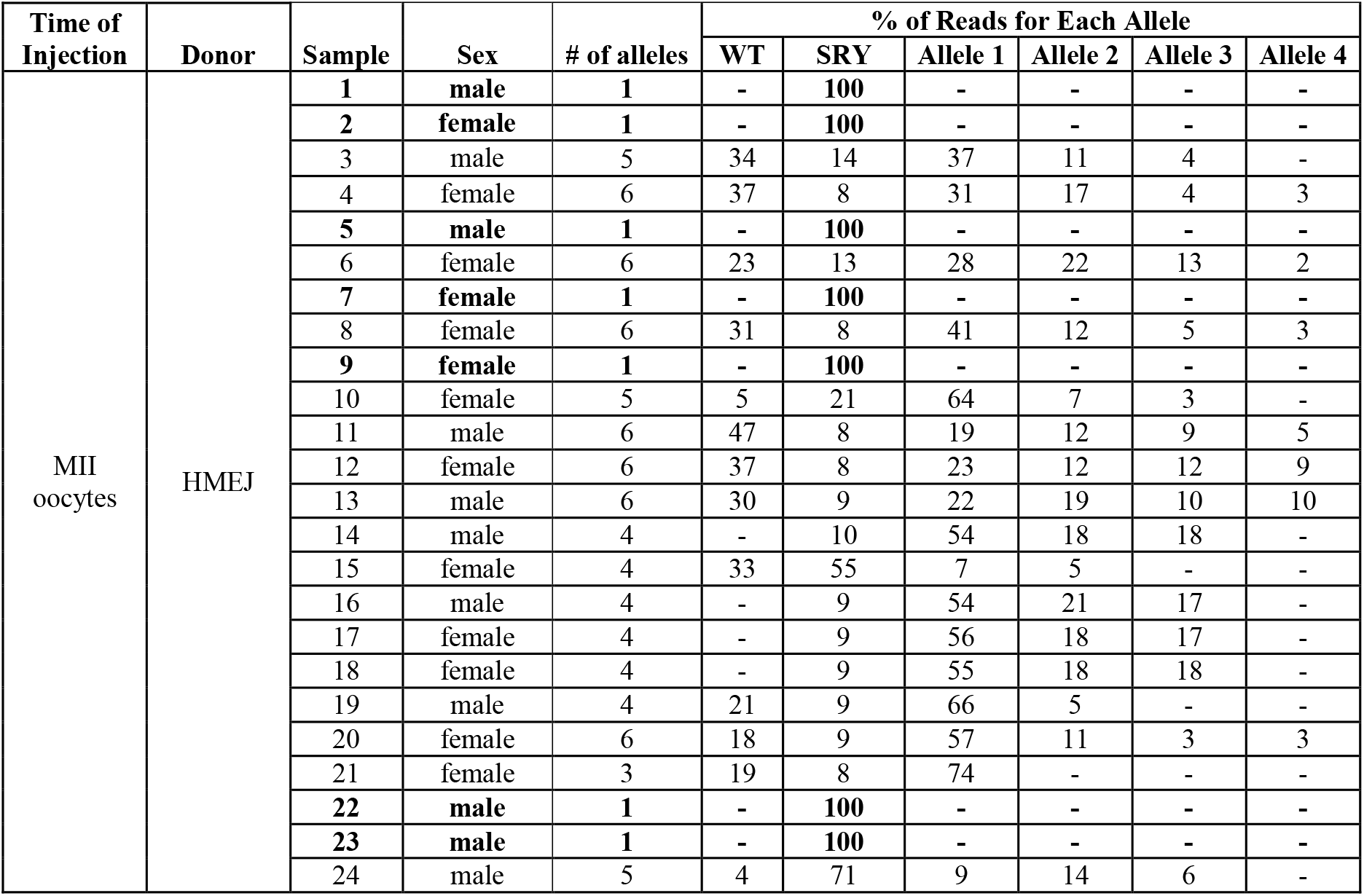
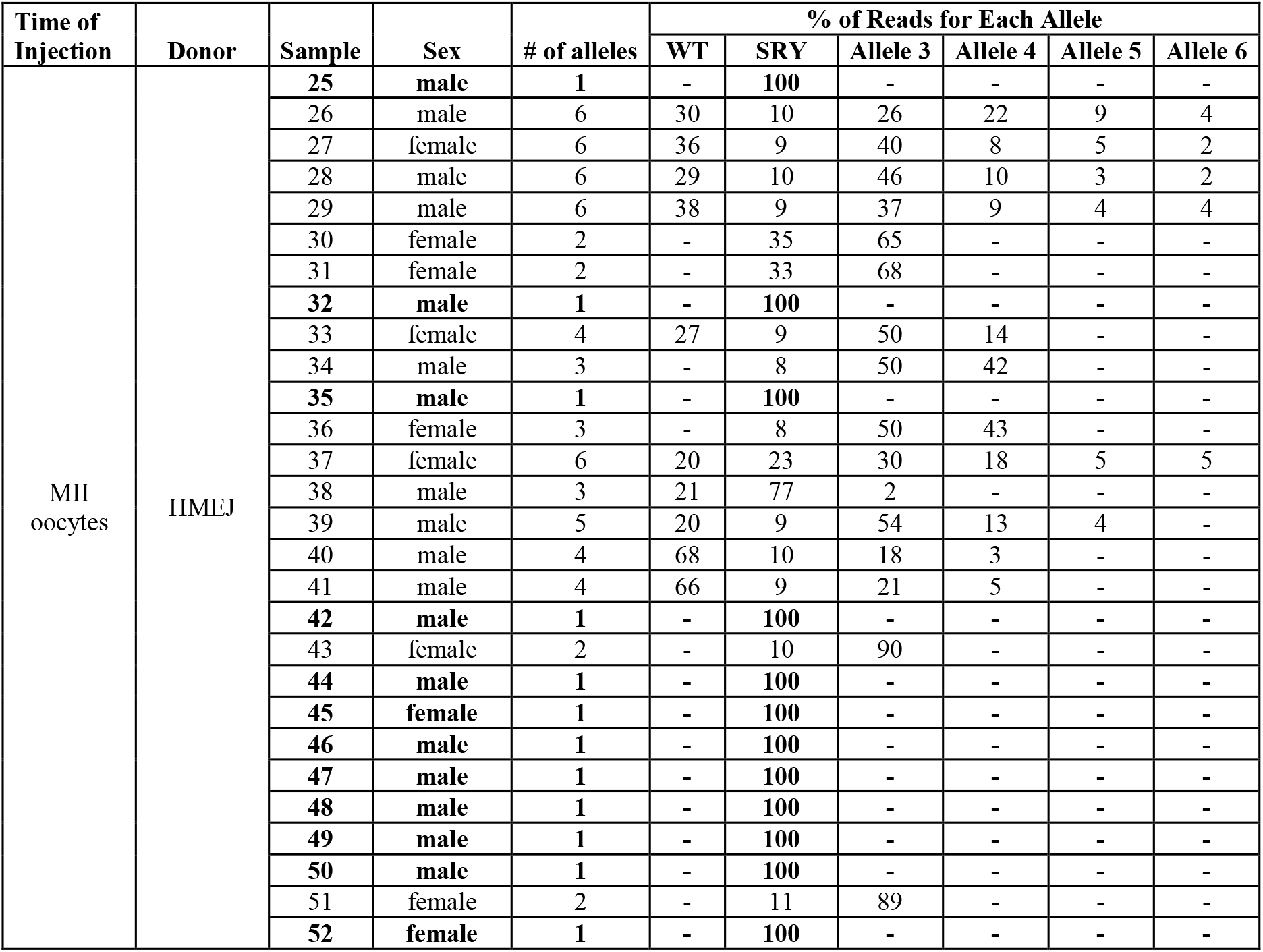
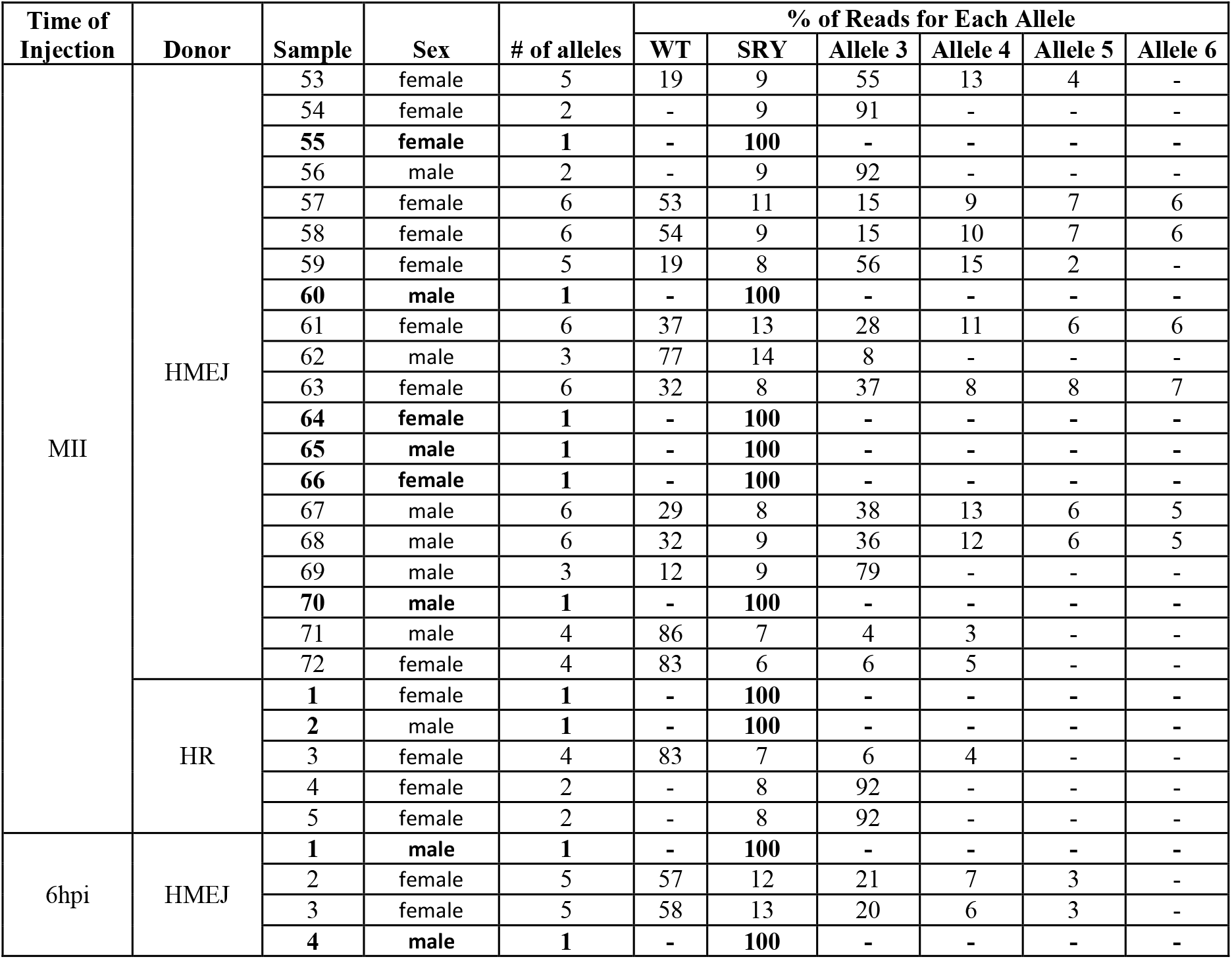
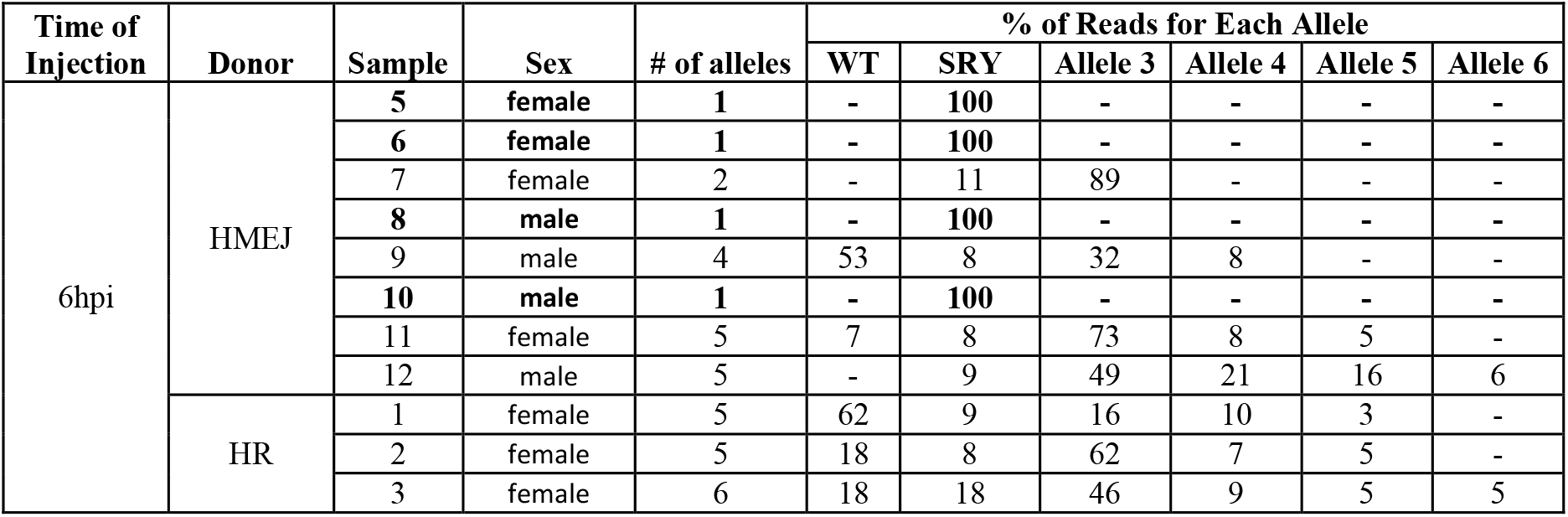
Number of alleles and percentage of each corresponding allele per sample detected at the cut-site of Cas9 protein injected embryos. WT = percentage of reads that were wild type sequence. SRY = insertion of the sex-determining region Y gene into the target location. Alleles 1-4 are percent reads with each of the alleles containing insertion or deletions found in the samples. Bold samples were non-mosaic and contained no wild type sequence.

**Supplementary Table S3.** Results from embryo transfer of MII injected oocytes followed by *in vitro* fertilization. No embryos were biopsied or transferred for trial one due to fungal contamination. Twelve of the 18 transferred blasts from trial four were vitrified, while six were fresh day eight blastocysts.

| Trial | Date | Biopsied | Vitrified | Blasts |  |
| --- | --- | --- | --- | --- | --- |
|  |  |  |  | Transferred | Pregnancy |
| 1 | 9/28/2018 | yes | no | 10 | 0 |
| 2 | 12/21/2018 | yes | yes | 12 | 0 |
|  |  |  | no | 6 | 0 |

**Supplementary Table S4.**
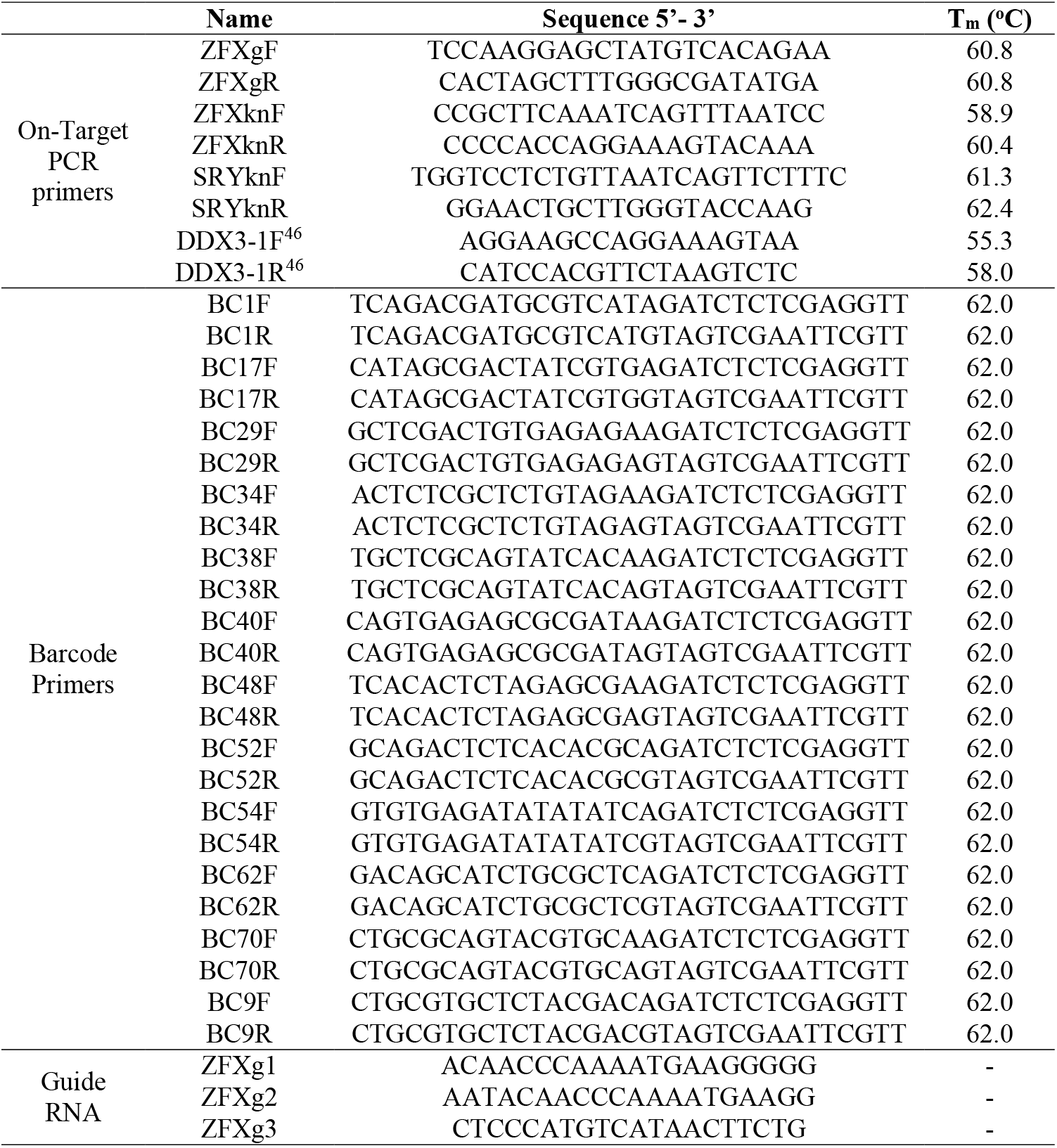
Sequence of primers used for PCR amplification of target region, evaluation of *SRY* knock-in, barcodes used for sequencing and guide-RNA sequences.

**Supplementary Table S5.** Development results from each collection by donor vector injected (hmejSRYp or hrSRYp), time of injection (MII oocytes or 6 hours post insemination (6hpi)), sex of the blastocysts, number of blastocysts collected, total number of embryos injected or cultured and resulting percentage of embryos that developed to the blastocyst stage.

| Collection | Donor | Time of Injection | Sex | Blast (n) | Total Embryos | Blast (%) |
| --- | --- | --- | --- | --- | --- | --- |
| 1 | control | control | control | 25 | 80 | 31.3 |
| 1 | HMEJ | MII | male | 2 | 20 | 10.0 |
| 2 | control | control | control | 24 | 80 | 30.0 |
| 2 | HMEJ | MII | female | 2 | 20 | 10.0 |
| 2 | HMEJ | MII | male | 2 | 20 | 10.0 |
| 3 | control | control | control | 26 | 80 | 32.5 |
| 3 | HMEJ | MII | female | 1 | 10 | 10.0 |
| 3 | HMEJ | MII | male | 1 | 10 | 10.0 |
| 4 | control | control | control | 23 | 80 | 38.8 |
| 4 | HMEJ | MII | male | 3 | 25 | 12.0 |
| 5 | control | control | control | 26 | 80 | 32.5 |
| 5 | HMEJ | MII | female | 7 | 100 | 7.0 |
| 5 | HMEJ | MII | male | 13 | 100 | 13.0 |
| 6 | control | control | control | 22 | 80 | 27.5 |
| 6 | HMEJ | MII | female | 5 | 50 | 10.0 |
| 6 | HMEJ | MII | male | 8 | 50 | 16.0 |
| 7 | control | control | control | 23 | 80 | 28.8 |
| 7 | HMEJ | MII | female | 12 | 100 | 12.0 |
| 7 | HMEJ | MII | male | 9 | 100 | 9.0 |
| 8 | control | control | control | 27 | 80 | 33.8 |
| 8 | HMEJ | MII | female | 2 | 20 | 10.0 |
| 8 | HMEJ | MII | male | 3 | 20 | 15.0 |
| 9 | control | control | control | 19 | 80 | 23.8 |
| 9 | HMEJ | MII | female | 3 | 100 | 3.0 |
| 9 | HMEJ | MII | male | 16 | 100 | 16.0 |
| 10 | control | control | control | 24 | 80 | 30.0 |
| 10 | HMEJ | MII | female | 9 | 40 | 22.5 |
| 10 | HMEJ | MII | male | 1 | 40 | 2.5 |
| 11 | control | control | control | 23 | 80 | 28.8 |
| 11 | HMEJ | MII | female | 13 | 100 | 13.0 |
| 11 | HMEJ | MII | male | 7 | 100 | 7.0 |
| 12 | control | control | control | 24 | 80 | 30.0 |
| 12 | HMEJ | MII | female | 5 | 50 | 10.0 |
| 12 | HMEJ | MII | male | 7 | 50 | 14.0 |
| 13 | control | control | control | 25 | 80 | 31.3 |
| 13 | HMEJ | MII | female | 2 | 40 | 5.0 |
| 13 | HMEJ | MII | male | 7 | 40 | 17.5 |
| 14 | control | control | control | 22 | 80 | 27.5 |
| 14 | HMEJ | MII | female | 4 | 100 | 4.0 |
| 14 | HMEJ | MII | male | 15 | 100 | 15.0 |
| 15 | control | control | control | 29 | 80 | 36.3 |
| 15 | HR | MII | female | 3 | 25 | 12.0 |
| 15 | HR | MII | male | 4 | 25 | 16.0 |
| 16 | control | control | control | 26 | 80 | 32.5 |
| 16 | HR | MII | female | 5 | 40 | 12.5 |
| 16 | HR | MII | male | 3 | 40 | 7.5 |
| 17 | control | control | control | 20 | 80 | 25.0 |
| 17 | HR | MII | female | 2 | 30 | 6.7 |
| 17 | HR | MII | male | 6 | 30 | 20.0 |
| 18 | control | control | control | 22 | 80 | 27.5 |
| 18 | HR | MII | female | 5 | 30 | 16.7 |
| 18 | HMEJ | 6hpi | female | 2 | 15 | 13.3 |
| 18 | HR | 6hpi | female | 5 | 15 | 33.3 |
| 18 | HR | MII | male | 3 | 30 | 10.0 |
| 18 | HMEJ | 6hpi | male | 2 | 15 | 13.3 |
| 18 | HR | 6hpi | male | 2 | 15 | 13.3 |
| 19 | control | control | control | 21 | 80 | 26.3 |
| 19 | HMEJ | 6hpi | female | 3 | 20 | 15.0 |
| 19 | HR | 6hpi | female | 1 | 20 | 5.0 |
| 19 | HMEJ | 6hpi | male | 6 | 20 | 30.0 |
| 19 | HR | 6hpi | male | 6 | 20 | 30.0 |
| 20 | control | control | control | 24 | 80 | 30.0 |
| 20 | HR | 6hpi | female | 5 | 30 | 16.7 |
| 21 | control | control | control | 23 | 80 | 28.8 |
| 21 | HR | 6hpi | female | 3 | 15 | 20.0 |
| 21 | HR | 6hpi | male | 3 | 15 | 20.0 |
| 22 | control | control | control | 22 | 80 | 27.5 |
| 22 | HR | 6hpi | female | 2 | 15 | 13.3 |
| 22 | HR | 6hpi | male | 1 | 15 | 6.7 |
| 23 | control | control | control | 24 | 80 | 30.0 |
| 23 | HMEJ | MII | male | 3 | 30 | 10.0 |
| 24 | control | control | control | 23 | 80 | 28.8 |
| 24 | HMEJ | MII | female | 4 | 25 | 16.0 |
| 24 | HMEJ | MII | male | 2 | 25 | 8.0 |
| 25 | control | control | control | 22 | 80 | 27.5 |
| 25 | HMEJ | MII | female | 1 | 30 | 3.3 |
| 25 | HMEJ | MII | male | 6 | 30 | 20.0 |
| 26 | control | control | control | 21 | 80 | 26.3 |
| 26 | HMEJ | 6hpi | female | 8 | 50 | 16.0 |
| 26 | HMEJ | 6hpi | male | 9 | 50 | 18.0 |
| 27 | control | control | control | 20 | 80 | 25.0 |
| 27 | HMEJ | MII | female | 4 | 30 | 13.3 |
| 27 | HMEJ | MII | male | 3 | 30 | 10.0 |
| 28 | control | control | control | 21 | 80 | 26.3 |
| 28 | HMEJ | MII | female | 1 | 15 | 6.7 |
| 28 | HMEJ | MII | male | 1 | 15 | 6.7 |
| 29 | control | control | control | 27 | 80 | 33.8 |
| 29 | HMEJ | MII | female | 11 | 100 | 11.0 |
| 29 | HMEJ | MII | male | 8 | 100 | 8.0 |
| 30 | control | control | control | 24 | 80 | 30.0 |
| 30 | HMEJ | MII | female | 3 | 30 | 10.0 |

**Supplementary Table S6.** Results for single blastocysts analyzed from each collected by donor vector injected (hmejSRYp or hrSRYp), time of injection (MII oocytes or 6 hours post insemination (6hpi), sex of the embryo, mutation in wild type allele, knock-in of SRY into the target location, type of knock-in (hemizygous, homozygous, homology independent target insertion (HITI) or mosaic) and the barcodes used for PacBio sequencing if sample was positive for SRY knock-in. fail = samples that had no sexing PCR product were excluded from further analysis.

| Collection | Sample | Donor | Time of Injection | Sex | Mutation | Knock-In | Knock-In Type | Forward Barcode | Reverse Barcode |
| --- | --- | --- | --- | --- | --- | --- | --- | --- | --- |
| 1 | 1 | HMEJ | MII | male | yes | no | - | - | - |
| 1 | 2 | HMEJ | MII | male | yes | no | - | - | - |
| 2 | 1 | HMEJ | MII | male | yes | yes | hemi | BC1F | BC1R |
| 2 | 2 | HMEJ | MII | male | no | no | - | - | - |
| 2 | 3 | HMEJ | MII | female | yes | yes | homo | BC1F | BC17R |
| 2 | 4 | HMEJ | MII | female | yes | no | - | - | - |
| 3 | 1 | HMEJ | MII | male | no | no | - | - | - |
| 3 | 2 | HMEJ | MII | female | yes | no | - | - | - |
| 4 | 1 | HMEJ | MII | male | yes | yes | mosaic | BC1F | BC29R |
| 4 | 2 | HMEJ | MII | male | no | no | - | - | - |
| 4 | 3 | HMEJ | MII | male | yes | no | - | - | - |
| 5 | 1 | HMEJ | MII | female | yes | yes | mosaic | BC1F | BC34R |
| 5 | 2 | HMEJ | MII | male | yes | no | - | - | - |
| 5 | 3 | HMEJ | MII | male | yes | no | - | - | - |
| 6 | 1 | HMEJ | MII | female | yes | no | - | - | - |
| 6 | 2 | HMEJ | MII | male | yes | yes | HITI | BC1F | BC38R |
| 6 | 3 | HMEJ | MII | female | yes | yes | mosaic | BC1F | BC40R |
| 6 | 4 | HMEJ | MII | male | yes | no | - | - | - |
| 7 | 1 | HMEJ | MII | male | no | no | - | - | - |
| 7 | 2 | HMEJ | MII | female | yes | no | - | - | - |
| 7 | 3 | HMEJ | MII | male | no | no | - | - | - |
| 7 | 4 | HMEJ | MII | female | no | no | - | - | - |
| 7 | 5 | HMEJ | MII | female | yes | yes | homo | BC1F | BC48R |
| 7 | 6 | HMEJ | MII | female | yes | yes | mosaic | BC1F | BC52R |
| 8 | 1 | HMEJ | MII | female | yes | yes | homo | BC1F | BC54R |
| 8 | 2 | HMEJ | MII | fail | - | - | - | - | - |
| 8 | 3 | HMEJ | MII | female | yes | yes | mosaic | BC1F | BC62R |
| 8 | 4 | HMEJ | MII | male | no | no | - | - | - |
| 8 | 5 | HMEJ | MII | male | yes | no | - | - | - |
| 8 | 6 | HMEJ | MII | male | no | no | - | - | - |
| 8 | 7 | HMEJ | MII | female | yes | no | - | - | - |
| 8 | 8 | HMEJ | MII | female | no | no | - | - | - |
| 9 | 1 | HMEJ | MII | fail | - | - | - | - | - |
| 9 | 2 | HMEJ | MII | fail | - | - | - | - | - |
| 9 | 3 | HMEJ | MII | male | yes | yes | mosaic | BC1F | BC70R |
| 9 | 4 | HMEJ | MII | fail | - | - | - | - | - |
| 9 | 5 | HMEJ | MII | fail | - | - | - | - | - |
| 9 | 6 | HMEJ | MII | fail | - | - | - | - | - |
| 9 | 7 | HMEJ | MII | female | yes | no | - | - | - |
| 9 | 8 | HMEJ | MII | fail | - | - | - | - | - |
| 10 | 1 | HMEJ | MII | male | no | no | - | - | - |
| 10 | 2 | HMEJ | MII | male | yes | no | - | - | - |
| 10 | 3 | HMEJ | MII | male | yes | no | - | - | - |
| 11 | 1 | HMEJ | MII | female | yes | yes | mosaic | BC1F | BC9R |
| 11 | 2 | HMEJ | MII | male | yes | no | - | - | - |
| 11 | 3 | HMEJ | MII | male | yes | no | - | - | - |
| 11 | 4 | HMEJ | MII | male | yes | yes | mosaic | BC17F | BC1R |
| 11 | 5 | HMEJ | MII | male | yes | no | - | - | - |
| 11 | 6 | HMEJ | MII | male | yes | yes | mosaic | BC17F | BC17R |
| 11 | 7 | HMEJ | MII | male | no | no | - | - | - |
| 12 | 1 | HMEJ | MII | female | yes | no | - | - | - |
| 12 | 2 | HMEJ | MII | fail | - | - | - | - | - |
| 12 | 3 | HMEJ | MII | female | no | no | - | - | - |
| 12 | 4 | HMEJ | MII | male | yes | no | - | - | - |
| 12 | 5 | HMEJ | MII | male | no | no | - | - | - |
| 12 | 6 | HMEJ | MII | female | yes | no | - | - | - |
| 12 | 7 | HMEJ | MII | male | yes | no | - | - | - |
| 12 | 8 | HMEJ | MII | male | no | no | - | - | - |
| 12 | 9 | HMEJ | MII | male | no | no | - | - | - |
| 12 | 10 | HMEJ | MII | female | no | no | - | - | - |
| 12 | 11 | HMEJ | MII | female | yes | yes | mosaic | BC17F | BC29R |
| 12 | 12 | HMEJ | MII | male | yes | yes | mosaic | BC17F | BC34R |
| 12 | 13 | HMEJ | MII | female | yes | yes | mosaic | BC17F | BC38R |
| 12 | 14 | HMEJ | MII | female | no | no | - | - | - |
| 12 | 15 | HMEJ | MII | female | yes | yes | mosaic | BC17F | BC40R |
| 13 | 1 | HMEJ | MII | male | yes | yes | mosaic | BC17F | BC48R |
| 13 | 2 | HMEJ | MII | female | yes | yes | mosaic | BC17F | BC52R |
| 13 | 3 | HMEJ | MII | female | yes | yes | mosaic | BC17F | BC54R |
| 13 | 4 | HMEJ | MII | female | yes | no | - | - | - |
| 13 | 5 | HMEJ | MII | male | yes | no | - | - | - |
| 13 | 6 | HMEJ | MII | male | no | no | - | - | - |
| 13 | 7 | HMEJ | MII | male | yes | yes | hemi | BC17F | BC62R |
| 13 | 8 | HMEJ | MII | male | yes | yes | hemi | BC17F | BC70R |
| 13 | 9 | HMEJ | MII | male | yes | yes | mosaic | BC17F | BC9R |
| 13 | 10 | HMEJ | MII | male | yes | yes | hemi | BC29F | BC1R |
| 13 | 11 | HMEJ | MII | male | yes | yes | mosaic | BC29F | BC17R |
| 13 | 12 | HMEJ | MII | female | yes | yes | mosaic | BC29F | BC29R |
| 13 | 13 | HMEJ | MII | male | yes | yes | mosaic | BC29F | BC34R |
| 13 | 14 | HMEJ | MII | female | yes | no | - | - | - |
| 13 | 15 | HMEJ | MII | fail | - | - | - | - | - |
| 13 | 16 | HMEJ | MII | female | yes | no | - | - | - |
| 13 | 17 | HMEJ | MII | male | yes | no | - | - | - |
| 13 | 18 | HMEJ | MII | male | yes | yes | mosaic | BC29F | BC38R |
| 14 | 1 | HMEJ | MII | fail | - | - | - | - | - |
| 14 | 2 | HMEJ | MII | fail | - | - | - | - | - |
| 14 | 3 | HMEJ | MII | fail | - | - | - | - | - |
| 14 | 4 | HMEJ | MII | female | no | no | - | - | - |
| 14 | 5 | HMEJ | MII | fail | - | - | - | - | - |
| 14 | 6 | HMEJ | MII | female | yes | no | - | - | - |
| 14 | 7 | HMEJ | MII | fail | - | - | - | - | - |
| 14 | 8 | HMEJ | MII | female | no | no | - | - | - |
| 14 | 9 | HMEJ | MII | fail | - | - | - | - | - |
| 15 | 1 | HMEJ | MII | male | no | no | - | - | - |
| 15 | 2 | HMEJ | MII | female | yes | yes | mosaic | BC29F | BC40R |
| 15 | 3 | HMEJ | MII | male | yes | no | - | - | - |
| 15 | 4 | HMEJ | MII | male | no | no | - | - | - |
| 15 | 5 | HMEJ | MII | female | yes | no | - | - | - |
| 15 | 6 | HMEJ | MII | male | yes | no | - | - | - |
| 15 | 7 | HMEJ | MII | male | yes | yes | mosaic | BC29F | BC48R |
| 15 | 8 | HMEJ | MII | female | yes | no | - | - | - |
| 15 | 9 | HMEJ | MII | male | no | no | - | - | - |
| 16 | 1 | HMEJ | MII | fail | - | - | - | - | - |
| 16 | 2 | HMEJ | MII | female | yes | no | - | - | - |
| 16 | 3 | HMEJ | MII | fail | - | - | - | - | - |
| 16 | 4 | HMEJ | MII | male | yes | yes | HITI | BC29F | BC52R |
| 16 | 5 | HMEJ | MII | female | yes | no | - | - | - |
| 16 | 6 | HMEJ | MII | fail | - | - | - | - | - |
| 16 | 7 | HMEJ | MII | female | yes | no | - | - | - |
| 16 | 8 | HMEJ | MII | female | yes | no | - | - | - |
| 16 | 9 | HMEJ | MII | male | yes | no | - | - | - |
| 16 | 10 | HMEJ | MII | male | no | no | - | - | - |
| 16 | 11 | HMEJ | MII | female | yes | yes | mosaic | BC29F | BC54R |
| 16 | 12 | HMEJ | MII | male | yes | no | - | - | - |
| 16 | 13 | HMEJ | MII | female | yes | no | - | - | - |
| 16 | 14 | HMEJ | MII | male | yes | yes | mosaic | BC29F | BC62R |
| 16 | 15 | HMEJ | MII | female | yes | no | - | - | - |
| 16 | 16 | HMEJ | MII | female | yes | no | - | - | - |
| 16 | 17 | HMEJ | MII | female | yes | no | - | - | - |
| 16 | 18 | HMEJ | MII | male | yes | yes | hemi | BC29F | BC70R |
| 16 | 19 | HMEJ | MII | female | yes | yes | mosaic | BC29F | BC9R |
| 16 | 20 | HMEJ | MII | female | yes | yes | mosaic | BC34F | BC1R |
| 16 | 21 | HMEJ | MII | male | yes | yes | mosaic | BC34F | BC17R |
| 16 | 22 | HMEJ | MII | male | yes | yes | mosaic | BC34F | BC29R |
| 16 | 23 | HMEJ | MII | female | yes | no | - | - | - |
| 16 | 24 | HMEJ | MII | male | yes | no | - | - | - |
| 17 | 1 | HMEJ | MII | female | yes | no | - | - | - |
| 17 | 2 | HMEJ | MII | male | yes | yes | mosaic | BC34F | BC34R |
| 17 | 3 | HMEJ | MII | male | yes | no | - | - | - |
| 17 | 4 | HMEJ | MII | female | yes | no | - | - | - |
| 17 | 5 | HMEJ | MII | male | no | no | - | - | - |
| 18 | 1 | HMEJ | MII | male | yes | no | - | - | - |
| 18 | 2 | HMEJ | MII | fail | - | - | - | - | - |
| 18 | 3 | HMEJ | MII | male | yes | yes | mosaic | BC34F | BC38R |
| 18 | 4 | HMEJ | MII | male | no | no | - | - | - |
| 18 | 5 | HMEJ | MII | male | yes | no | - | - | - |
| 18 | 6 | HMEJ | MII | fail | - | - | - | - | - |
| 18 | 7 | HMEJ | MII | male | yes | yes | hemi | BC34F | BC40R |
| 18 | 8 | HMEJ | MII | female | yes | yes | mosaic | BC34F | BC48R |
| 18 | 9 | HMEJ | MII | male | yes | yes | HITI | BC34F | BC52R |
| 18 | 10 | HMEJ | MII | female | yes | yes | HITI | BC34F | BC54R |
| 18 | 11 | HMEJ | MII | male | yes | no | - | - | - |
| 18 | 12 | HMEJ | MII | male | no | no | - | - | - |
| 18 | 13 | HMEJ | MII | male | yes | yes | HITI | BC34F | BC62R |
| 18 | 14 | HMEJ | MII | male | yes | no | - | - | - |
| 18 | 15 | HMEJ | MII | male | yes | no | - | - | - |
| 18 | 16 | HMEJ | MII | male | yes | yes | HITI | BC34F | BC70R |
| 18 | 17 | HMEJ | MII | male | yes | no | - | - | - |
| 18 | 18 | HMEJ | MII | male | yes | yes | HITI | BC34F | BC9R |
| 18 | 19 | HMEJ | MII | male | yes | yes | HITI | BC38F | BC1R |
| 18 | 20 | HMEJ | MII | male | yes | yes | HITI | BC38F | BC17R |
| 19 | 1 | HMEJ | MII | male | yes | no | - | - | - |
| 19 | 2 | HMEJ | MII | female | yes | yes | mosaic | BC38F | BC29R |
| 19 | 3 | HMEJ | MII | female | yes | yes | HITI | BC38F | BC34R |
| 19 | 4 | HMEJ | MII | fail | - | - | - | - | - |
| 19 | 5 | HMEJ | MII | female | yes | yes | mosaic | BC38F | BC38R |
| 19 | 6 | HMEJ | MII | female | yes | yes | mosaic | BC38F | BC40R |
| 19 | 7 | HMEJ | MII | female | yes | yes | homo | BC38F | BC48R |
| 19 | 8 | HMEJ | MII | fail | - | - | - | - | - |
| 19 | 9 | HMEJ | MII | female | yes | yes | mosaic | BC38F | BC52R |
| 19 | 10 | HMEJ | MII | female | yes | yes | mosaic | BC38F | BC54R |
| 19 | 11 | HMEJ | MII | female | no | no | - | - | - |
| 19 | 12 | HMEJ | MII | female | no | no | - | - | - |
| 20 | 1 | HMEJ | MII | female | yes | yes | mosaic | BC38F | BC62R |
| 20 | 2 | HMEJ | MII | male | yes | no | - | - | - |
| 20 | 3 | HMEJ | MII | male | no | no | - | - | - |
| 20 | 4 | HMEJ | MII | male | yes | no | - | - | - |
| 20 | 5 | HMEJ | MII | female | yes | yes | mosaic | BC38F | BC70R |
| 20 | 6 | HMEJ | MII | female | yes | no | - | - | - |
| 20 | 7 | HMEJ | MII | male | yes | yes | hemi | BC38F | BC9R |
| 20 | 8 | HMEJ | MII | female | yes | no | - | - | - |
| 20 | 9 | HMEJ | MII | male | yes | no | - | - | - |
| 20 | 10 | HMEJ | MII | female | yes | yes | mosaic | BC40F | BC1R |
| 20 | 11 | HMEJ | MII | male | no | no | - | - | - |
| 20 | 12 | HMEJ | MII | female | yes | no | - | - | - |
| 20 | 13 | HMEJ | MII | female | yes | no | - | - | - |
| 20 | 14 | HMEJ | MII | female | no | no | - | - | - |
| 20 | 15 | HMEJ | MII | male | yes | yes | mosaic | BC40F | BC17R |
| 20 | 16 | HMEJ | MII | female | yes | yes | mosaic | BC40F | BC29R |
| 20 | 17 | HMEJ | MII | female | yes | no | - | - | - |
| 21 | 1 | HMEJ | MII | female | yes | yes | homo | BC40F | BC34R |
| 21 | 2 | HMEJ | MII | male | yes | yes | HITI | BC40F | BC38R |
| 21 | 3 | HMEJ | MII | male | no | no | - | - | - |
| 21 | 4 | HMEJ | MII | male | yes | no | - | - | - |
| 21 | 5 | HMEJ | MII | female | yes | yes | homo | BC40F | BC40R |
| 21 | 6 | HMEJ | MII | male | yes | yes | mosaic | BC40F | BC48R |
| 21 | 7 | HMEJ | MII | female | no | no | - | - | - |
| 21 | 8 | HMEJ | MII | female | yes | no | - | - | - |
| 21 | 9 | HMEJ | MII | male | yes | no | - | - | - |
| 21 | 10 | HMEJ | MII | female | no | no | - | - | - |
| 21 | 11 | HMEJ | MII | male | yes | no | - | - | - |
| 21 | 12 | HMEJ | MII | male | yes | no | - | - | - |
| 22 | 1 | HMEJ | MII | female | no | no | - | - | - |
| 22 | 2 | HMEJ | MII | male | yes | no | - | - | - |
| 22 | 3 | HMEJ | MII | male | no | no | - | - | - |
| 22 | 4 | HMEJ | MII | female | yes | no | - | - | - |
| 22 | 5 | HMEJ | MII | male | yes | yes | mosaic | BC40F | BC52R |
| 22 | 6 | HMEJ | MII | fail | - | - | - | - | - |
| 22 | 7 | HMEJ | MII | male | no | no | - | - | - |
| 22 | 8 | HMEJ | MII | male | yes | no | - | - | - |
| 22 | 9 | HMEJ | MII | male | yes | no | - | - | - |
| 22 | 9 | HMEJ | MII | male | yes | no | - | - | - |
| 23 | 1 | HMEJ | MII | male | yes | no | - | - | - |
| 23 | 2 | HMEJ | MII | male | yes | no | - | - | - |
| 23 | 3 | HMEJ | MII | male | no | no | - | - | - |
| 23 | 4 | HMEJ | MII | fail | - | - | - | - | - |
| 23 | 5 | HMEJ | MII | male | no | no | - | - | - |
| 23 | 6 | HMEJ | MII | female | yes | no | - | - | - |
| 23 | 7 | HMEJ | MII | male | yes | no | - | - | - |
| 23 | 8 | HMEJ | MII | male | yes | yes | mosaic | BC40F | BC54R |
| 23 | 9 | HMEJ | MII | male | yes | no | - | - | - |
| 23 | 10 | HMEJ | MII | male | yes | no | - | - | - |
| 23 | 11 | HMEJ | MII | male | yes | no | - | - | - |
| 23 | 12 | HMEJ | MII | female | yes | no | - | - | - |
| 23 | 13 | HMEJ | MII | male | yes | yes | hemi | BC40F | BC62R |
| 23 | 14 | HMEJ | MII | male | yes | no | - | - | - |
| 23 | 15 | HMEJ | MII | female | no | no | - | - | - |
| 23 | 16 | HMEJ | MII | male | yes | yes | mosaic | BC40F | BC70R |
| 23 | 17 | HMEJ | MII | male | yes | no | - | - | - |
| 23 | 18 | HMEJ | MII | female | yes | yes | mosaic | BC40F | BC9R |
| 23 | 19 | HMEJ | MII | male | no | no | - | - | - |
| 23 | 20 | HMEJ | MII | fail | - | - | - | - | - |
| 23 | 21 | HMEJ | MII | male | no | no | - | - | - |
| 24 | 1 | HMEJ | 6hpi | male | yes | yes | hemi | BC48F | BC1R |
| 24 | 2 | HMEJ | 6hpi | male | yes | no | - | - | - |
| 24 | 3 | HMEJ | 6hpi | male | yes | no | - | - | - |
| 24 | 4 | HMEJ | 6hpi | male | yes | no | - | - | - |
| 24 | 5 | HMEJ | 6hpi | female | yes | no | - | - | - |
| 24 | 6 | HMEJ | 6hpi | female | yes | no | - | - | - |
| 24 | 7 | HMEJ | 6hpi | female | yes | yes | mosaic | BC48F | BC17R |
| 24 | 8 | HMEJ | 6hpi | male | yes | no | - | - | - |
| 24 | 9 | HMEJ | 6hpi | female | yes | yes | mosaic | BC48F | BC29R |
| 24 | 10 | HMEJ | 6hpi | female | yes | no | - | - | - |
| 24 | 11 | HMEJ | 6hpi | male | yes | no | - | - | - |
| 24 | 12 | HMEJ | 6hpi | female | yes | no | - | - | - |
| 24 | 13 | HMEJ | 6hpi | female | no | no | - | - | - |
| 24 | 14 | HMEJ | 6hpi | male | yes | no | - | - | - |
| 24 | 15 | HMEJ | 6hpi | male | yes | no | - | - | - |
| 24 | 16 | HMEJ | 6hpi | fail | - | - | - | - | - |
| 24 | 17 | HMEJ | 6hpi | male | yes | yes | HITI | BC48F | BC34R |
| 24 | 18 | HMEJ | 6hpi | female | yes | yes | HITI | BC48F | BC38R |
| 25 | 1 | HR | MII | male | no | no | - | - | - |
| 25 | 2 | HR | MII | male | yes | no | - | - | - |
| 25 | 3 | HR | MII | female | yes | no | - | - | - |
| 25 | 4 | HR | MII | male | no | no | - | - | - |
| 25 | 5 | HR | MII | male | yes | no | - | - | - |
| 25 | 6 | HR | MII | female | yes | no | - | - | - |
| 25 | 7 | HR | MII | fail | - | - | - | - | - |
| 25 | 8 | HR | MII | female | no | no | - | - | - |
| 26 | 1 | HR | MII | female | yes | no | - | - | - |
| 26 | 2 | HR | MII | male | yes | no | - | - | - |
| 26 | 3 | HR | MII | female | yes | yes | mosaic | BC48F | BC40R |
| 26 | 4 | HR | MII | female | no | no | - | - | - |
| 26 | 5 | HR | MII | female | yes | no | - | - | - |
| 26 | 6 | HR | MII | male | yes | no | - | - | - |
| 26 | 7 | HR | MII | female | no | no | - | - | - |
| 26 | 8 | HR | MII | male | yes | no | - | - | - |
| 27 | 1 | HR | MII | female | yes | no | - | - | - |
| 27 | 2 | HR | MII | male | no | no | - | - | - |
| 27 | 3 | HR | MII | male | yes | no | - | - | - |
| 27 | 4 | HR | MII | male | yes | yes | mosaic | BC48F | BC48R |
| 27 | 5 | HR | MII | male | yes | no | - | - | - |
| 27 | 6 | HR | MII | male | yes | no | - | - | - |
| 27 | 7 | HR | MII | male | yes | no | - | - | - |
| 27 | 8 | HR | MII | female | yes | yes | homo | BC48F | BC52R |
| 28 | 1 | HR | MII | female | yes | no | - | - | - |
| 28 | 2 | HR | MII | female | yes | no | - | - | - |
| 28 | 3 | HR | MII | male | yes | no | - | - | - |
| 28 | 4 | HR | MII | female | yes | yes | homo | BC48F | BC54R |
| 28 | 5 | HR | MII | male | yes | no | - | - | - |
| 28 | 6 | HR | MII | female | yes | yes | mosaic | BC48F | BC62R |
| 28 | 7 | HR | MII | male | yes | no | - | - | - |
| 29 | 1 | HR | 6hpi | female | yes | yes | mosaic | BC48F | BC70R |
| 29 | 2 | HR | 6hpi | female | yes | yes | mosaic | BC48F | BC9R |
| 29 | 3 | HR | 6hpi | male | yes | no | - | - | - |
| 29 | 4 | HR | 6hpi | female | yes | yes | mosaic | BC52F | BC1R |
| 29 | 5 | HR | 6hpi | female | yes | no | - | - | - |
| 29 | 6 | HR | 6hpi | female | yes | no | - | - | - |
| 29 | 7 | HR | 6hpi | male | yes | no | - | - | - |
| 30 | 1 | HMEJ | 6hpi | female | yes | yes | homo | BC52F | BC17R |
| 30 | 2 | HMEJ | 6hpi | male | yes | no | - | - | - |
| 30 | 3 | HMEJ | 6hpi | female | yes | yes | mosaic | BC52F | BC29R |
| 30 | 4 | HMEJ | 6hpi | male | yes | no | - | - | - |
| 31 | 1 | HMEJ | 6hpi | male | yes | yes | HITI | BC52F | BC34R |
| 31 | 2 | HMEJ | 6hpi | male | yes | yes | mosaic | BC52F | BC38R |
| 31 | 3 | HMEJ | 6hpi | female | yes | no | - | - | - |
| 31 | 4 | HMEJ | 6hpi | male | yes | yes | HITI | BC52F | BC40R |
| 31 | 5 | HMEJ | 6hpi | female | yes | yes | mosaic | BC52F | BC48R |
| 31 | 6 | HMEJ | 6hpi | male | yes | no | - | - | - |
| 31 | 7 | HMEJ | 6hpi | male | yes | yes | mosaic | BC52F | BC52R |
| 31 | 8 | HMEJ | 6hpi | male | no | no | - | - | - |
| 31 | 9 | HMEJ | 6hpi | female | yes | no | - | - | - |
| 32 | 1 | HR | 6hpi | male | yes | no | - | - | - |
| 32 | 2 | HR | 6hpi | female | yes | no | - | - | - |
| 32 | 3 | HR | 6hpi | male | yes | no | - | - | - |
| 32 | 4 | HR | 6hpi | male | no | no | - | - | - |
| 32 | 5 | HR | 6hpi | male | yes | no | - | - | - |
| 32 | 6 | HR | 6hpi | male | yes | no | - | - | - |
| 32 | 7 | HR | 6hpi | male | yes | no | - | - | - |
| 33 | 1 | HR | 6hpi | female | yes | no | - | - | - |
| 33 | 2 | HR | 6hpi | fail | - | - | - | - | - |
| 33 | 3 | HR | 6hpi | female | no | no | - | - | - |
| 33 | 4 | HR | 6hpi | fail | - | - | - | - | - |
| 33 | 5 | HR | 6hpi | fail | - | - | - | - | - |
| 33 | 6 | HR | 6hpi | female | yes | no | - | - | - |
| 33 | 7 | HR | 6hpi | female | yes | no | - | - | - |
| 33 | 8 | HR | 6hpi | female | yes | no | - | - | - |
| 34 | 1 | HR | 6hpi | female | yes | no | - | - | - |
| 34 | 2 | HR | 6hpi | male | yes | no | - | - | - |
| 34 | 3 | HR | 6hpi | female | yes | no | - | - | - |
| 34 | 4 | HR | 6hpi | female | yes | no | - | - | - |
| 34 | 5 | HR | 6hpi | male | yes | no | - | - | - |
| 34 | 6 | HR | 6hpi | male | no | no | - | - | - |
| 35 | 1 | HR | 6hpi | male | yes | no | - | - | - |
| 35 | 2 | HR | 6hpi | female | no | no | - | - | - |
| 35 | 3 | HR | 6hpi | female | yes | no | - | - | - |

**Supplementary Figure S1.**
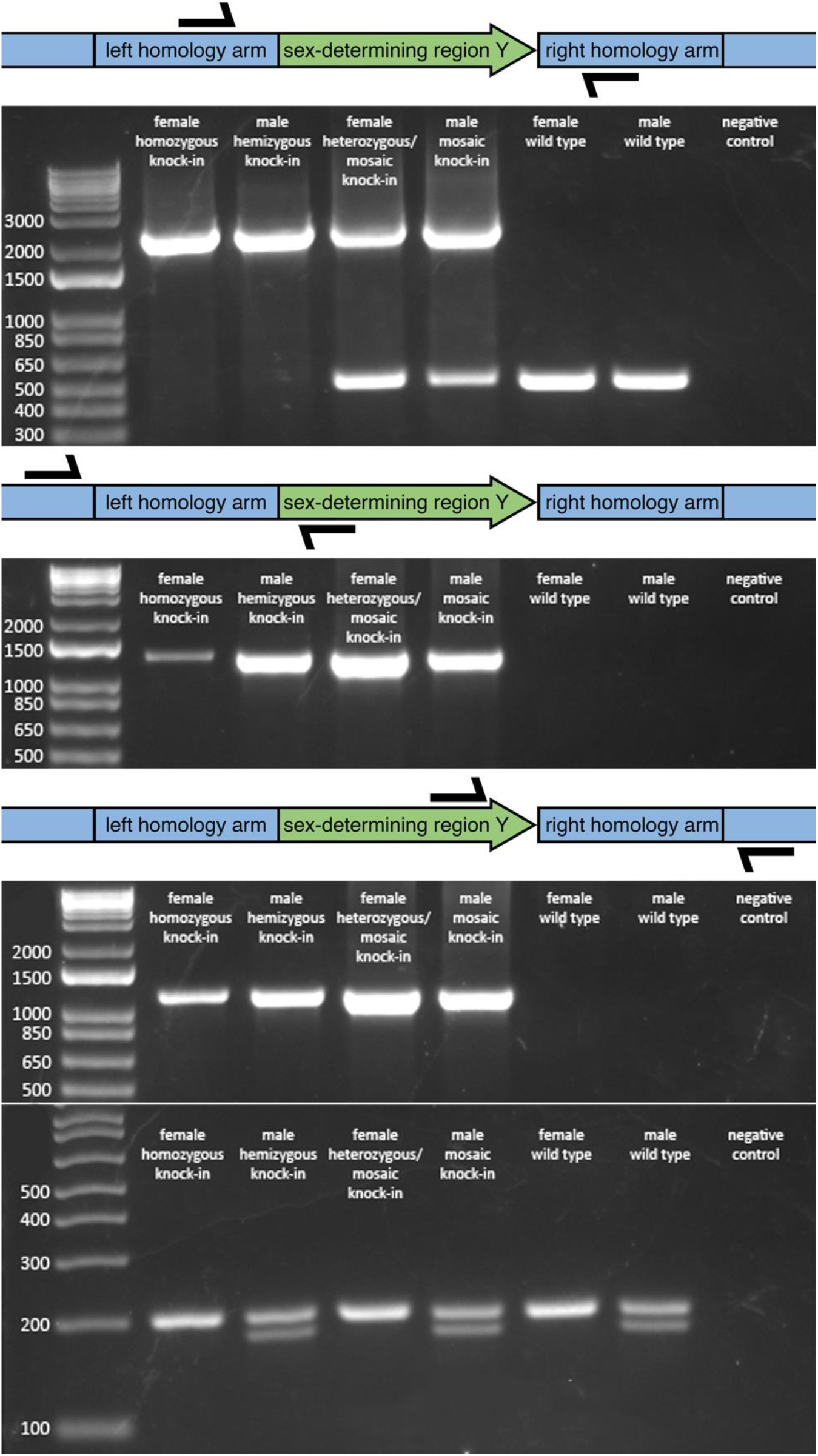
PCR analysis of embryos for *SRY* knock-in at the ZFX locus. (a) Expected wild type product 520bp, knock-in product 2349bp. (b) Expected *SRY* knock-in product 1427bp. (c) Expected *SRY* knock-in product 1307bp. (d): female expected product 208bp, male expected product 189 and 208bp.

**Supplementary Figure S2.**
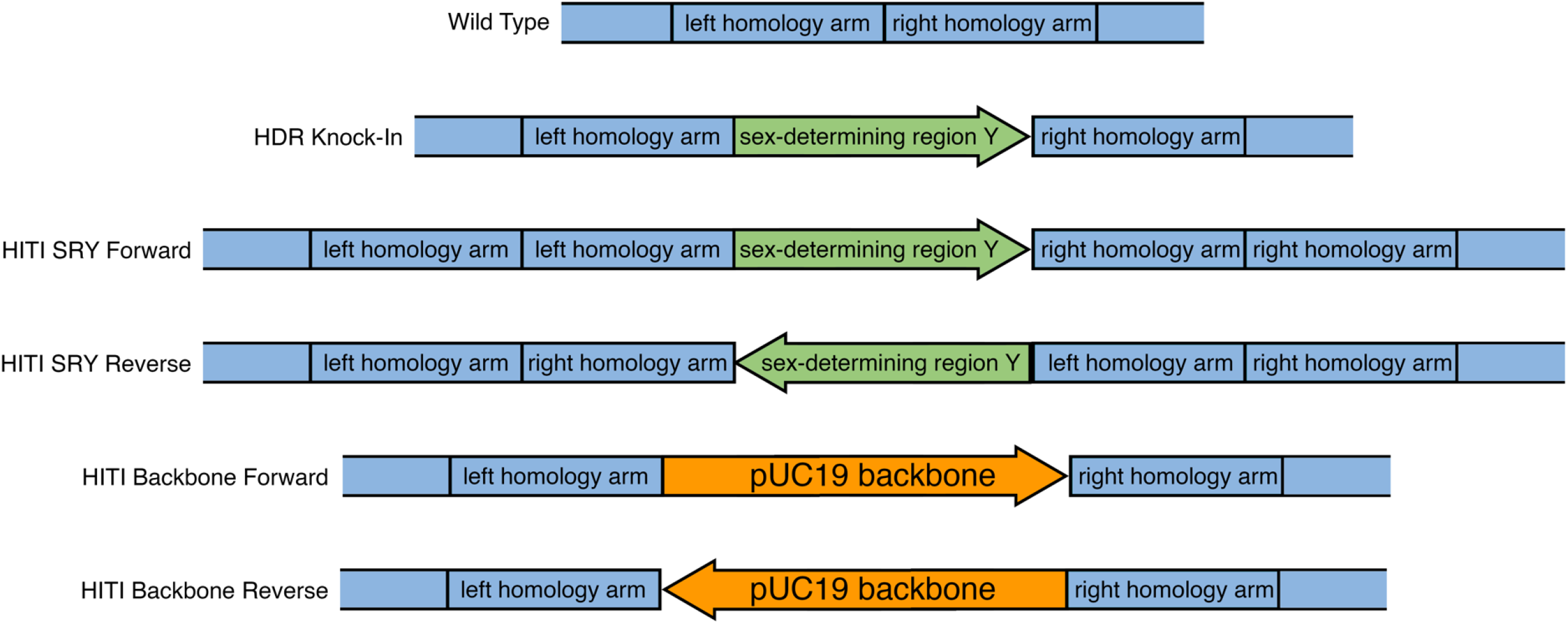
Schematic representation of target sequences used to map PacBio reads against for evaluating presence of wildtype and knock-in of the sex-determining region Y (*SRY*) or plasmid backbone by homology directed repair (HDR) or homology independent targeted repair (HITI).

## REFERENCES

1 Van Eenennaam, A. L. Genetic modification of food animals. Curr Opin Biotechnol 44, 27–34, doi:10.1016/j.copbio.2016.10.007 (2017).

2 Tait-Burkard, C. et al. Livestock 2.0 - genome editing for fitter, healthier, and more productive farmed animals. Genome Biol 19, 204, doi:10.1186/s13059-018-1583-1 (2018).

3 Bishop, T. F. & Van Eenennaam, A. L. Genome editing approaches to augment livestock breeding programs. J Exp Biol 223, doi:10.1242/jeb.207159 (2020).

4 Georges, M., Charlier, C. & Hayes, B. Harnessing genomic information for livestock improvement. Nat Rev Genet 20, 135–156, doi:10.1038/s41576-018-0082-2 (2019).

5 Tan, W., Proudfoot, C., Lillico, S. G. & Whitelaw, C. B. Gene targeting, genome editing: from Dolly to editors. Transgenic Res 25, 273–287, doi:10.1007/s11248-016-9932-x (2016).

6 Yum, S. Y., Youn, K. Y., Choi, W. J. & Jang, G. Development of genome engineering technologies in cattle: from random to specific. J Anim Sci Biotechnol 9, 16, doi:10.1186/s40104-018-0232-6 (2018).

7 Mclean, Z., Oback, B. & Laible, G. Embryo-mediated genome editing for accelerated genetic improvement of livestock. Frontiers of Agricultural Science and Engineering, doi:doi: 10.15302/J-FASE-2019305 (2020).

8 Mehravar, M., Shirazi, A., Nazari, M. & Banan, M. Mosaicism in CRISPR/Cas9-mediated genome editing. Dev Biol 445, 156–162, doi:10.1016/j.ydbio.2018.10.008 (2019).

9 Hennig, S. L. et al. Evaluation of mosaicism and off-target mutations in CRISPR-mediated genome edited bovine embryos. bioRxiv 2020.2006.2004.134759, doi:10.1101/2020.06.04.134759 (2020).

10 Lamas-Toranzo, I. et al. Strategies to reduce genetic mosaicism following CRISPR-mediated genome edition in bovine embryos. Scientific Reports 9, 14900, doi:10.1038/s41598-019-51366-8 (2019).

11 Danner, E. et al. Control of gene editing by manipulation of DNA repair mechanisms. Mamm Genome 28, 262–274, doi:10.1007/s00335-017-9688-5 (2017).

12 Hustedt, N. & Durocher, D. The control of DNA repair by the cell cycle. Nat Cell Biol 19, 1–9, doi:10.1038/ncb3452 (2016).

13 Yao, X. et al. Homology-mediated end joining-based targeted integration using CRISPR/Cas9. Cell Res 27, 801–814, doi:10.1038/cr.2017.76 (2017).

14 Sfeir, A. & Symington, L. S. Microhomology-Mediated End Joining: A Back-up Survival Mechanism or Dedicated Pathway? Trends Biochem Sci 40, 701–714, doi:10.1016/j.tibs.2015.08.006 (2015).

15 Suzuki, K. et al. In vivo genome editing via CRISPR/Cas9 mediated homology-independent targeted integration. Nature 540, 144–149, doi:10.1038/nature20565 (2016).

16 Ma, T. et al. An AANAT/ASMT transgenic animal model constructed with CRISPR/Cas9 system serving as the mammary gland bioreactor to produce melatonin-enriched milk in sheep. J Pineal Res 63, doi:10.1111/jpi.12406 (2017).

17 Lillico, S. G. et al. Mammalian interspecies substitution of immune modulatory alleles by genome editing. Sci Rep 6, 21645, doi:10.1038/srep21645 (2016).

18 Wei, J. et al. Efficient introgression of allelic variants by embryo-mediated editing of the bovine genome. Sci Rep 5, 11735, doi:10.1038/srep11735 (2015).

19 Wei, J. et al. Cattle with a precise, zygote-mediated deletion safely eliminate the major milk allergen beta-lactoglobulin. Sci Rep 8, 7661, doi:10.1038/s41598-018-25654-8 (2018).

20 Navarro-Serna, S., Vilarino, M., Park, I., Gadea, J. & Ross, P. J. Livestock gene editing by one-step embryo manipulation. Journal of Equine Veterinary Science, 103025, doi:https://doi.org/10.1016/j.jevs.2020.103025 (2020).

21 Daigneault, B. W., Rajput, S., Smith, G. W. & Ross, P. J. Embryonic POU5F1 is Required for Expanded Bovine Blastocyst Formation. Sci Rep 8, 7753, doi:10.1038/s41598-018-25964-x (2018).

22 Vilarino, M. et al. CRISPR/Cas9 microinjection in oocytes disables pancreas development in sheep. Sci Rep 7, 17472, doi:10.1038/s41598-017-17805-0 (2017).

23 Tanihara, F. et al. Effects of concentration of CRISPR/Cas9 components on genetic mosaicism in cytoplasmic microinjected porcine embryos. Journal of Reproduction and Development 65, 209–214 (2019).

24 Miao, D., Giassetti, M. I., Ciccarelli, M., Lopez-Biladeau, B. & Oatley, J. M. Simplified pipelines for genetic engineering of mammalian embryos by CRISPR-Cas9 electroporationdagger. Biol Reprod 101, 177–187, doi:10.1093/biolre/ioz075 (2019).

25 Namula, Z. et al. Genome mutation after the introduction of the gene editing by electroporation of Cas9 protein (GEEP) system into bovine putative zygotes. In Vitro Cell Dev Biol Anim 55, 598–603, doi:10.1007/s11626-019-00385-w (2019).

26 Brophy, B. et al. in Proceedings of the World Congress on Genetics Applied to Livestock Production Vol. Technologies - Gene Editing 1118 (2018).

27 Chen, S., Lee, B., Lee, A. Y. F., Modzelewski, A. J. & He, L. Highly Efficient Mouse Genome Editing by CRISPR Ribonucleoprotein Electroporation of Zygotes. Journal of Biological Chemistry 291, 14457–14467, doi:10.1074/jbc.M116.733154 (2016).

28 Okamoto, S., Amaishi, Y., Maki, I., Enoki, T. & Mineno, J. Highly efficient genome editing for single-base substitutions using optimized ssODNs with Cas9-RNPs. Scientific Reports 9, 4811, doi:10.1038/s41598-019-41121-4 (2019).

29 Yoshimi, K. et al. ssODN-mediated knock-in with CRISPR-Cas for large genomic regions in zygotes. Nat Commun 7, 10431, doi:10.1038/ncomms10431 (2016).

30 McFarlane, G. R., Salvesen, H. A., Sternberg, A. & Lillico, S. G. On-Farm Livestock Genome Editing Using Cutting Edge Reproductive Technologies. Frontiers in Sustainable Food Systems 3, doi:10.3389/fsufs.2019.00106 (2019).

31 Yoon, Y. et al. Streamlined ex vivo and in vivo genome editing in mouse embryos using recombinant adeno-associated viruses. Nature communications 9, 1–12 (2018).

32 Chen, S. et al. CRISPR-READI: Efficient Generation of Knockin Mice by CRISPR RNP Electroporation and AAV Donor Infection. Cell Rep 27, 3780–3789 e3784, doi:10.1016/j.celrep.2019.05.103 (2019).

33 Yoon, Y. et al. Streamlined ex vivo and in vivo genome editing in mouse embryos using recombinant adeno-associated viruses. Nature Communications 9, 412, doi:10.1038/s41467-017-02706-7 (2018).

34 de Sousa, R. V. et al. Biopsy of bovine embryos produced in vivo and in vitro does not affect pregnancy rates. Theriogenology 90, 25–31, doi:10.1016/j.theriogenology.2016.11.003 (2017).

35 Vilarino, M. et al. Mosaicism diminishes the value of pre-implantation embryo biopsies for detecting CRISPR/Cas9 induced mutations in sheep. Transgenic Res 27, 525–537, doi:10.1007/s11248-018-0094-x (2018).

36 Bakhtari, A. & Ross, P. J. DPPA3 prevents cytosine hydroxymethylation of the maternal pronucleus and is required for normal development in bovine embryos. Epigenetics 9, 1271–1279, doi:10.4161/epi.32087 (2014).

37 Bogliotti, Y. S., Vilarino, M. & Ross, P. J. Laser-assisted Cytoplasmic Microinjection in Livestock Zygotes. J Vis Exp, doi:10.3791/54465 (2016).

38 Koressaar, T. & Remm, M. Enhancements and modifications of primer design program Primer3. Bioinformatics 23, 1289–1291, doi:10.1093/bioinformatics/btm091 (2007).

39 Untergasser, A. et al. Primer3--new capabilities and interfaces. Nucleic Acids Res 40, e115, doi:10.1093/nar/gks596 (2012).

40 Daneau, I., Houde, A., Ethier, J. F., Lussier, J. G. & Silversides, D. W. Bovine SRY gene locus: cloning and testicular expression. Biology of Reproduction 52, 591–599 (1995).

41 Langmead, B. & Salzberg, S. L. Fast gapped-read alignment with Bowtie 2. Nat Methods 9, 357–359, doi:10.1038/nmeth.1923 (2012).

42 Robinson, J. T., Thorvaldsdottir, H., Wenger, A. M., Zehir, A. & Mesirov, J. P. Variant Review with the Integrative Genomics Viewer. Cancer Res 77, e31–e34, doi:10.1158/0008-5472.CAN-17-0337 (2017).

43 Li, H. Aligning sequence reads, clone sequences and assembly contigs with BWA-MEM. arXiv 1303.3997v (2013).

44 Li, H. et al. The Sequence Alignment/Map format and SAMtools. Bioinformatics 25, 2078–2079, doi:10.1093/bioinformatics/btp352 (2009).

45 Lindsay, H. et al. CrispRVariants charts the mutation spectrum of genome engineering experiments. Nat Biotechnol 34, 701–702, doi:10.1038/nbt.3628 (2016).

46 Gokulakrishnan, P., Kumar, R. R., Sharma, B. D., Mendiratta, S. K. & Sharma, D. Sex Determination of Cattle Meat by Polymerase Chain Reaction Amplification of the DEAD Box Protein (DDX3X/DDX3Y) Gene. Asian-Australas J Anim Sci 25, 733–737, doi:10.5713/ajas.2012.12003 (2012).

